# Cancer blues? A validated judgment bias task suggests pessimism in nude mice with tumors

**DOI:** 10.1101/2021.02.13.431089

**Authors:** A. Resasco, A. MacLellan, M. A. Ayala, L. Kitchenham, A. M. Edwards, S. Lam, S. Dejardin, G. Mason

## Abstract

In humans, affective states can bias responses to ambiguous information: a phenomenon termed judgment bias (JB). Judgment biases have great potential for assessing affective states in animals, in both animal welfare and biomedical research. New animal JB tasks require construct validation, but for laboratory mice (*Mus musculus*), the most common research vertebrate, a valid JB task has proved elusive. Here (Experiment 1), we demonstrate construct validity for a novel mouse JB test: an olfactory Go/Go task in which subjects dig for high- or low-value food rewards. In C57BL/6 and Balb/c mice faced with ambiguous cues, latencies to dig were sensitive to high/low welfare housing, environmentally-enriched animals responding with relative ‘optimism’ through shorter latencies. Illustrating the versatility of a validated JB task across fields of research, it further allowed us to test hypotheses about the mood-altering effects of cancer (Experiment 2). Male nude mice bearing subcutaneous lung adenocarcinomas responded more pessimistically than healthy controls to ambiguous cues. Similar effects were not seen in females, however. To our knowledge, this is the first validation of a mouse JB task and the first demonstration of pessimism in tumor-bearing animals. This task, especially if refined to improve its sensitivity, thus has great potential for investigating mouse welfare, the links between affective state and disease, depression-like states in animals, and hypotheses regarding the neurobiological mechanisms that underlie affect-mediated biases in judgment.

## 1. Introduction

Of the 115+ million animals used annually in biomedical research [1], most are rodents. They are often used to model potentially distressing conditions like cancer, arthritis and psychiatric disorders (e.g. anxiety, depression). But even conventional practices like handling (e.g. [2]) and the use of small, non-enriched cages (e.g. [3–5]) can compromise their wellbeing. These welfare costs can modify experimental outcomes in undesired directions [6]. They also have ethical implications, especially given the poor replicability [7] and translatability of biomedical research [8–10]. Our focus here is a potential method for assessing affective states (emotions and long-term moods [11]) in mice: the most widely used vertebrate in both basic and translational research [12]. Such methods are important for assessing mouse welfare, and for understanding the neurobiological mechanisms underlying normal and pathological affective functioning.

In humans, affective states modulate the interpretation of ambiguous information, a phenomenon known as judgment bias (JB). JB refers to the way that individuals experiencing negative affect (e.g. anxiety, depression) can process ambiguous information (e.g. neutral facial expressions) ‘pessimistically’, as if negative, while individuals in positive states might demonstrate more ‘optimistic’ interpretations of the same ambiguous cues [see 13—15]. In animal JB studies, optimism can be operationalized as increased expectations of reward when faced with ambiguous cues, and pessimism, by increased expectations of punishment [16]. Harding et al., [17] pioneered this method of animal JB assessment: rats trained that one cue predicts reward while another predicts punishment, were exposed to ambiguous (intermediate) cues. Rats exposed to unpredictable housing showed pessimistic JBs, treating the ambiguous cues as if predicting punishment. Since this seminal work, JB tasks have gained popularity as potentially powerful tools for assessing animal affect due to their sensitivity to changes in both valence and intensity of these states [18]. Thus JB tasks have been developed for a wide range of species (e.g. dogs, sheep, horses, honeybees), using a variety of cues (e.g. visual, olfactory, tactile), and across diverse fields of research (e.g. behavioral biology, neuroscience and animal welfare) [19,20].

For mice, however, validated JB tasks had remained elusive. Valid JB tasks must meet two technical criteria: that animals discriminate between positive and negative cues, and then interpret intermediate cues as ambiguous [20,21]. But like any putative indicator of affective state, they must also demonstrate construct validity: sensitivity to deliberate affect manipulations (c.f. [22,23]). For mice, previous efforts have either not attempted construct validation (5/15 experiments [24–26]), or attempted it and failed (10/15 experiments [27–33]; Table S1). Here, we therefore aimed to validate a novel JB task, manipulating affective state through the use of highly preferred environmentally enriched cages [34], versus conventional cages known to induce stress [35], anxiety [36,37], and depression-like effects [35,38,39]. Environmental enrichment (modification of an animal’s environment to improve well-being and meet species-specific needs [40]), has been used in neuroscience for decades for its positive effects in neuroplasticity and disease recovery [41]. Morphological and physiological changes in the brain due to enrichment have also been associated with improved welfare [42], and JB has been shown to be sensitive to the effects of enrichment in other species (e.g. rats [43,44]).

In a second experiment, we applied this newly validated task to mice with tumors, to assess its utility in translational biomedical research. It is well established that cancer can be debilitating when tumors cause pain and discomfort (e.g. [45]), and rodent welfare guidelines for oncology already focus on such harms (e.g. [46]). However, tumors are known to reduce human well-being at much earlier stages: tumors can induce depression-like feelings of sadness and hopelessness [47,48], even before cancer is diagnosed (e.g. [49,50]), thanks to elevated pro-inflammatory cytokines [51,52]. Mice with tumors likewise show signs of depression (e.g. increased anhedonia [53]). And again these reflect inflammatory responses [54–57], and are manifest before clinical signs emerge [46,58]. However, these subtle changes have received negligible attention in mouse welfare guidelines. Nor have more nuanced measures of mood yet been developed for researchers interested in the translational benefits of mouse models of cancer. To bridge these research gaps, we thus aimed to assess mood in mice with tumors through judgment bias.

## 2. Materials, methods and results

### 2.1. Ethical note

Both experiments were approved by institutional ethics committees. Experiment 1 (AUP #3700) complied with Canadian Council on Animal Care guidelines, and Experiment 2 (protocol number 42-1-14T) complied with Guidelines for the Welfare and Use of Animals in Cancer Research [46]. One C57 was removed before testing for barbering a cagemate (Experiment 1), and one male nude mouse was removed due an eye abscess (Experiment 2). This report also meets ARRIVE (Animal Research: Reporting of *In Vivo* Experiments) requirements [59].

### 2.2. Experiment 1: Validating a Novel JB Task with Housing-Manipulated Affective States

#### 2.2.1. Animals and Housing

Eighteen C57BL/6NCrl (‘C57’) and 18 Balb/cAnNCrl (‘Balb’) females were purchased from Charles River (Raleigh, North Carolina) at 3-4 weeks old. Females were chosen to allow for the combined use of group housing (important since mice are a social species), and environmental enrichment without the risk of resource guarding aggression that can be problematic in male mice [60]. Mice were randomly assigned to open-top enriched or conventional housing treatments (respectively EH or CH). Here they lived in mixed strain groups (c.f. [61]), each cage containing one C57 and one Balb, plus two DBA/2NCrl cagemates used in another experiment. CH comprised transparent polyethylene laboratory cages (27L × 16W × 12H cm, Allentown Inc.; n = 9), with corn cob bedding (Envigo, Mississauga, Ontario, Canada), a paper cup and two types of nesting material (crinkled paper strips and cotton pads; Fig. 1 A). EH cages were large (60L × 60W × 30H cm, n=9), opaque plastic with one transparent red plastic window, containing a variety of enrichments that facilitate species-typical behaviors (e.g. hiding, climbing, chewing, and nesting [c.f. 39]; Fig. 1 B and C). Attached to each enriched cage was a standard ‘annex’ cage that mice could freely access via a tunnel. Mice were trained to enter the annex cage for a food treat when a cup full of sweet oat cereal (Cheerios) was shaken; the access tunnel could then be blocked allowing for ease of catching and handling in the annex. Handling for both treatments always followed cup or tunnel methods to minimize aversive effects [2]. The room was kept at 21±1°C and 35-55% relative humidity, on a reverse 12:12 hour light cycle (lights off at 06:00). Food (Harlan® Teklad, Global Diet 14% protein) and water were *ad libitum.*

**Fig 1.**
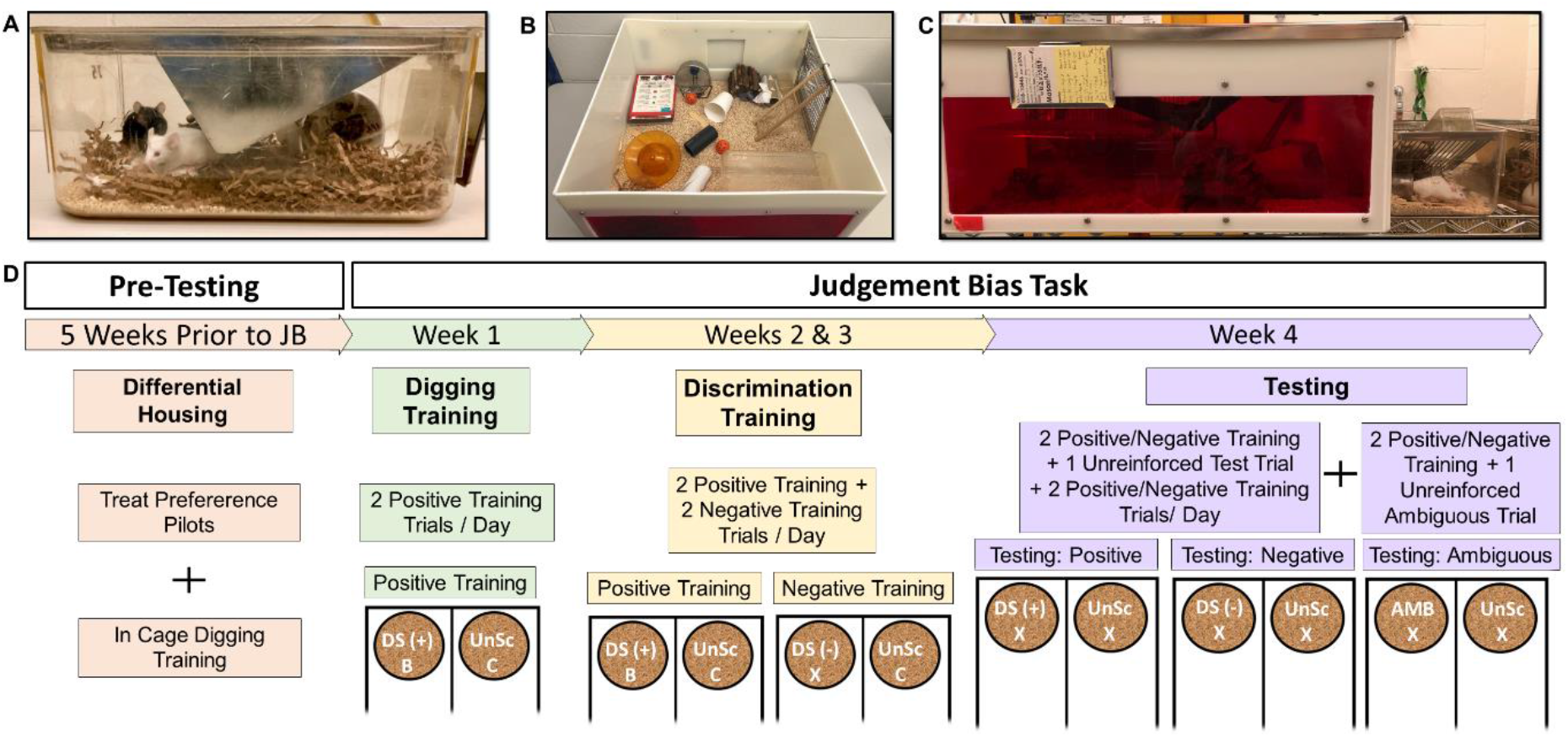
Housing treatments and timeline for Experiment 1. A. Conventional ‘shoebox’ laboratory cage with a paper cup and two types of nest material; B and C. Upper and front view of the enriched cage, respectively; a standard ‘annex’ cage was attached to one of the sides of the enriched cage to facilitate handling. D. Timeline and summary of positive, negative and ambiguous training and test trials for Experiment 1. DS(+): positive discriminative stimulus, DS(-): negative discriminative stimulus, AMB: ambiguous mixture (50% vanilla-50%mint), B: banana chip, C: Rodent diet (‘chow’), X: no food rewards.

#### 2.2.2. Judgment bias (JB) training and testing

Our olfactory, digging-based task utilized a “Go/Go” design that was divided into three phases (Fig. 1 D). All JB training and testing was conducted under red light in an experimental area of the colony room (separated by a plastic curtain), between 08:00 and 18:00. Mice were pseudorandomly assigned to an experimenter blind to treatment (AM or AR), counterbalancing across housing and strains. Randomization was conducted through an online random order generator [62], unless otherwise noted. Mice were fasted for one hour prior to training or testing throughout all phases to increase motivation for food, while also maintaining the preference for the high-over the low-value rewards [30]. After fasting, mice were moved from their home cage to a transport cage (27L × 16W × 12H cm, Allentown Inc) by a familiar research assistant and placed in the experimental area. The order in which mice were tested was random across days since EH mice were opportunistically caught in the annex cage (see above). Between trials, all plastic components of the apparatus were wiped thoroughly with 70% ethanol and disposable materials were replaced.

The apparatus (Fig. 2) comprised a start compartment and two arms, each containing a scent dispenser at its entrance (a cotton-filled tissue cassette) and a 6.5L × 6.5W × 4H cm corncob-filled pot at its far end. To prevent the scent of the buried treats from revealing which pot was rewarded, an inaccessible treat compartment was located at the bottom of each pot with perforated plastic to allow odor transmission. Treats included in the inaccessible compartment were dependent on which treat (if any) was accessible, so that each pot always included a total of one chow piece and one banana chip across compartments. The whole apparatus was topped with a transparent plexiglass cover.

**Fig 2.**
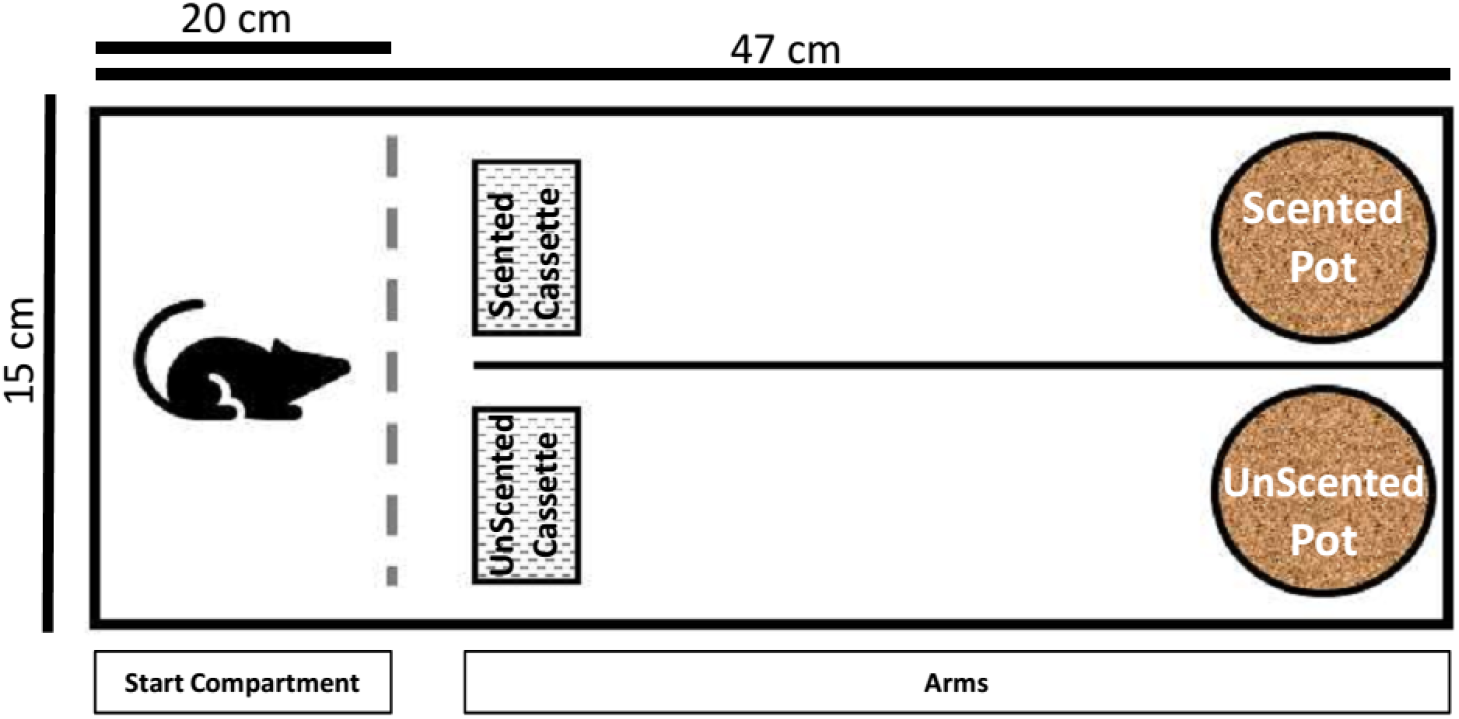
Judgment bias apparatus used in Experiments 1 and 2. The dotted line represents the sliding door that was opened at the beginning of each trial.

Pilot tests identified preferred treats (Desjardin unpubl); dried, sweetened banana chips (Stock and Barrel) were selected as the high-value reward and regular rodent chow was used as low-value reward. Vanilla and mint essences (Fleibor S.R.L, Buenos Aires, Argentina), diluted 1:4 in distilled water [63,64], acted as cues (discriminative stimuli: DS). In each trial, one arm of the apparatus was always unscented (marked with distilled water), predicting a buried low-value reward (rodent chow). DS+ or DS-solution (0.1 ml) was applied to the scent dispenser and corncob of the scented arm, respectively predicting buried high-value rewards, banana chips (in positive trials) or no reward (negative trials) (Table 1). Throughout all phases, to facilitate learning and prevent extinction, if a mouse was still eating a reward when the trial ended she was allowed 30 seconds to eat before being handled. Additionally, if she had not yet found the reward by the end of a trial, the appropriate treat was placed on top of the bedding and the mouse was gently guided to it (and given 30 seconds to eat). Each mouse’s DS+ (mint or vanilla) and the side of the scented arm (left or right) were pseudorandomly assigned, counterbalancing across strain, housing and experimenter.

**Table 1.**
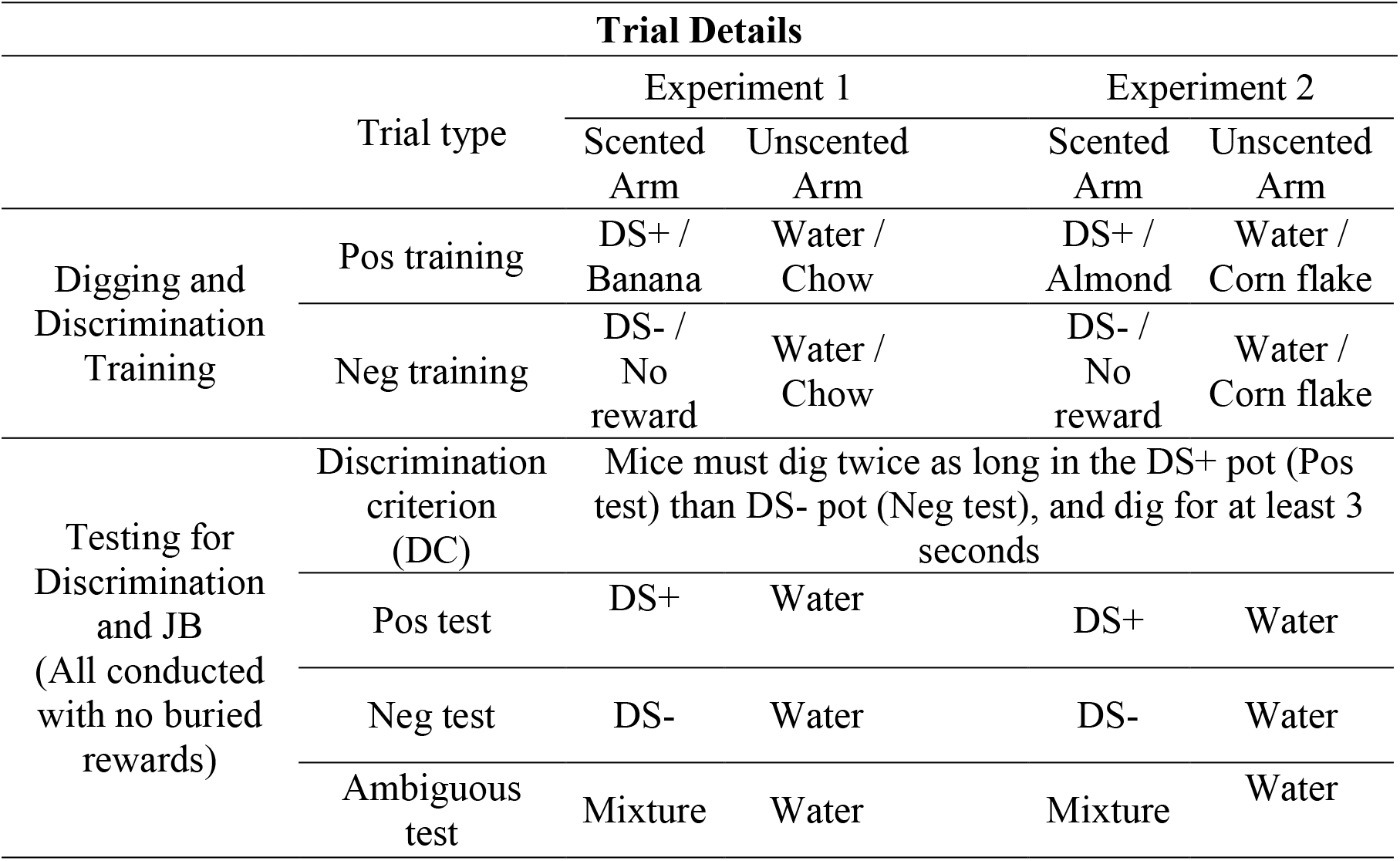
Summary of the trial details in Experiments 1 and 2. DS(+): positive discriminative stimulus, DS(-): negative discriminative stimulus, Pos: positive, Neg: negative. See Supplemental Table S2 for expanded table.

#### 2.2.3. Digging Training

One week before training (when 8-9 weeks old), mice were habituated to digging pots with two being placed in their cages daily for 10 minutes, one containing low-value rewards, the other high-value rewards (see Fig. 1 D for full experimental timeline). Digging training in the apparatus began the following week. Here, treats were placed on top of the corn cob bedding on Day 1 and progressively buried in the following four days until they were completely buried at the bottom of the pot by Day 5. Two positive trials were run per day, each lasting 5 minutes. Mice were allowed to freely explore the apparatus and their latency to start eating each reward was live recorded. Preferences for banana chips over rodent chow were confirmed on the final day (see Results). All mice were able to find the rewards by the end of this phase allowing them to move on to the next stage.

#### 2.2.4. Discrimination Training

This phase introduced the negative trial and lasted 10 days, with two trials per mouse each day. For Days 1-5, the first trial was positive and the second was negative. During Days 6-10, the order of the positive and negative trial was randomized daily. Latencies to dig and to eat in both arms were live scored.

#### 2.2.5. Testing for discrimination learning

To confirm successful discrimination of DS+ and DS-, mice underwent 4 reinforced trials daily for 2-4 days (the length of this phase being variable and determined by how quickly each mouse reached discrimination criterion; see below). These trials were divided into two blocks, with one unreinforced (test) trial in the middle to assess their responses to each DS (Table S2). The order of positive and negative reinforced trials before and after the test was randomized for each mouse. Positive and negative test trials were presented in alternating order across days (e.g. Day One DS+ test, Day 2 DS-test, Day 3 DS+ test, etc.). Test trials lasted 2 minutes and were videoed. Appropriate rewards were placed on the corncob after each trial (banana for DS+, chow for DS-), and mice were allowed 30 seconds to eat before being moved back to their transport cage.

Latency to dig, as well as the total duration of digging, in the first and the full 2 minutes of test trials were recorded by two observers blind to treatment, and their values were averaged (with videos showing marked discrepancies being re-scored). The discrimination criterion set required mice to dig for at least twice as long in the DS+ arm than in the DS-arm in the first minute of testing (with a minimum DS+ digging time of 3 seconds). Mice who met discrimination criteria moved on to ambiguous cue testing the following day (see below). Mice who did not yet meet criteria continued to be presented with unrewarded DS+ and DS-trials until criteria was met. Mice who did not meet criteria within 4 days were excluded from ambiguous trials (see results).

On the day of ambiguous testing, mice received one positive and one negative trial in random order, followed by a video-recorded ambiguous unreinforced trial in which an ambiguous mixture (50% diluted mint, 50% diluted vanilla) marked the scented arm. Again, videos were scored by two observers, blind to treatment, for latency to dig and digging duration.

#### 2.2.6. In-cage behavioral observations

Behavioral observations were conducted to check for expected differences in welfare between EH and CH mice (e.g. higher levels of stereotypic behaviors in the latter; [34,39]). Data were collected via live scan sampling during the dark, active phase. A silent observer scanned each cage every 15 minutes for four hours, starting two hours after lights off [c.f. 65]. The first observed behavior for each mouse was categorized according to the ethogram (Table S3). Since EH mice had more opportunities to be out of sight, each behavior was calculated as a proportion of visible scans.

#### 2.2.7. Statistical analyses for Experiment 1

Generalized Linear Mixed Models in SAS®9.4 were used, on data transformed where needed to meet assumptions (normality and homogeneity of residuals). Where assumptions could not be met, non-parametric tests were used instead (and noted in text). Treat preferences were confirmed during digging training by assessing latency to eat high- and low-value rewards. The repeated measures model therefore included Reward, Housing, Strain, DS+ Odor and all two way interactions, plus Cage (a random effect nested in Housing and DS+ odor) and Mouse ID (a random effect nested in Cage, Housing, DS+ odor and Strain). To test for judgment bias, repeated measures models were run to assess both latency to dig and digging duration in the scented arm for positive, negative and ambiguous test trials. Trial Type, Housing, Strain, Trial Type*Housing, DS+ odor, Trial Type*Strain, Trial Type*DS+ odor, Trial Type*Housing*DS+ odor, Cage (a random effect nested in Housing and DS+ odor) and Mouse ID (a random effect nested in Cage, Housing, DS+ odor and Strain) were always included in Experiment 1 models. To select which additional main and interactive effects to include (e.g. Tester ID and its interactions), a stepwise forward selection process using corrected Akaike’s Information Criteria (c.f. [66]) identified the most parsimonious final models. These were then run using maximum-likelihood estimations (the experiment becoming unbalanced when not all mice met discrimination criteria, c.f. [20]). Since Housing was the treatment of interest, simple effects were calculated from the Trial Type*Housing, using the SLICEDIFF command when calculating the Least Squares Means [67]. One-tailed Ps were used since only one specific effect would validate the task (c.f.[68]): shorter latencies and longer digging times for EH than CH mice, in ambiguous trials only. Finally, effect sizes (Cohen’s d) were calculated, and ANOVA tables were used to assess whether Treatment*Trial Type contributed significant variation. To confirm housing effects, levels of home cage stereotypic behavior (SB) and time spent inactive but awake (IBA) were assessed. For SB, terms included in the model were Housing, Strain, Housing*Strain and Cage (as a random effect nested within housing). Home cage IBA data did not meet assumptions of normality and homogeneity, so a Wilcoxon rank sum test was used instead.

### 2.3. Experiment 1 results

All but 4 C57 mice met discrimination criteria (n=31). Treat preference was confirmed by the lower latencies to eat the high-value reward over the low-value reward the last day of digging training (*F*_1,32_=80.46, *p*<0.0001, *Cohen’s d=*2.215). Housing*Reward (*F*_1,32_=4.12, *p*=0.005) and Strain*Reward were significant (*F*_1,32_=6.89, *p*=0.007), but banana was still preferred in all subgroups (p<0.0001). During tests for JB, Trial Type*DS+ odor was significant for both latency to dig and digging duration (respectively *F*_2,62_=5.74, *p*=0.005 and *F*_2,62_=18.88, *p*<0.0001), Mint DS+ mice unexpectedly treating intermediate odor mixtures as positive (as if 100% mint), but Vanilla DS+ mice treating intermediate odor mixtures as ambiguous as required for a valid JB task (Fig. 3 A, Table S4). There were also significant strain differences in latency (latency: *F*_1,31_=4.87, *p*=0.018, *Cohen’s d*=-0.799; duration: *F*_1,31_=3.82, *p*=0.030, *Cohen’s d*=0.484), C57s from both housing conditions showing shorter latencies and digging durations than Balbs across trials.

**Fig 3.**
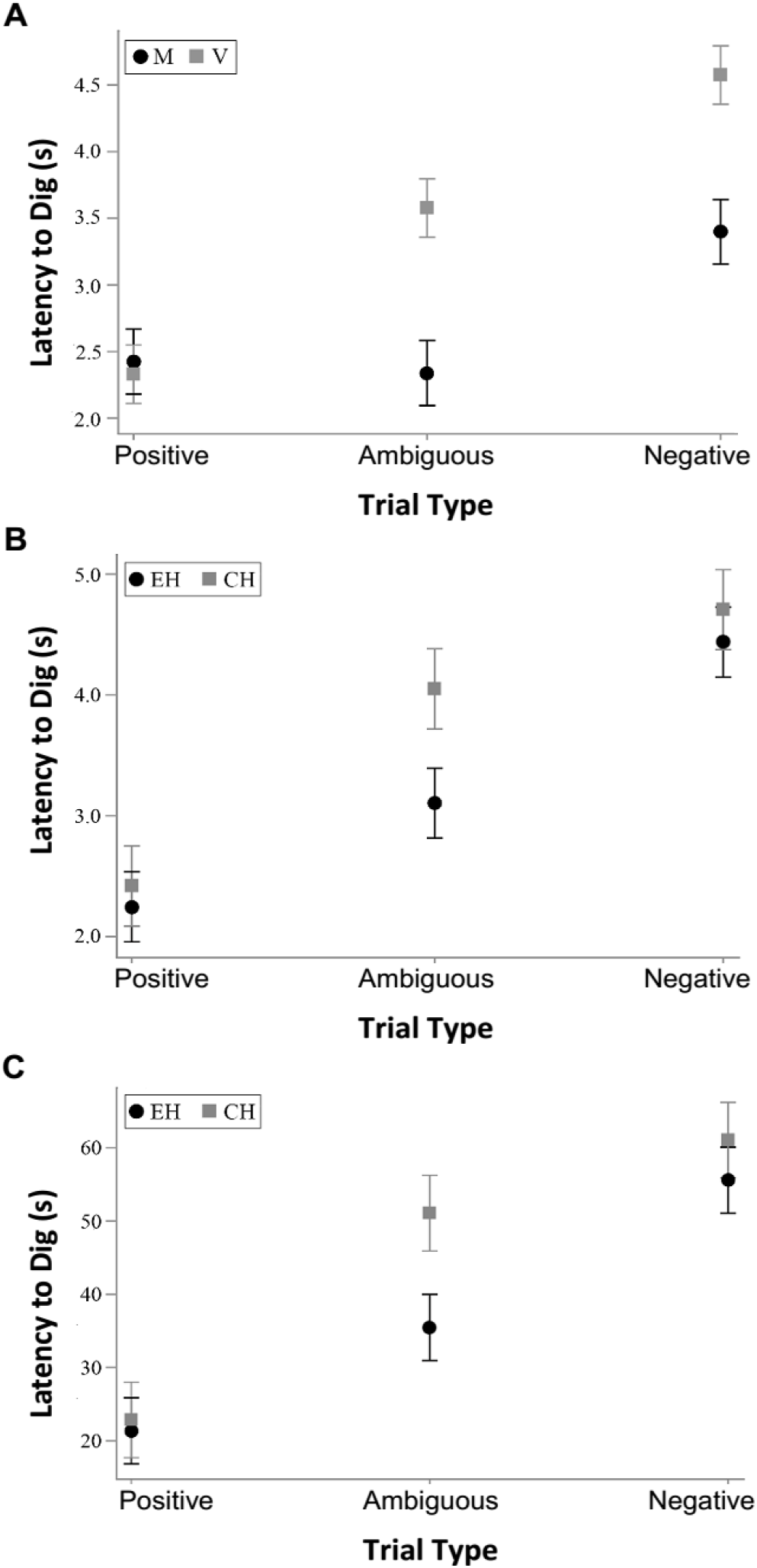
Digging latency least square means (± standard error) during positive, negative and ambiguous test trials. A. 2min digging latency in mice receiving mint (M, n=15) or vanilla (V, n=16) as the positive discriminative stimulus (DS+) (data logarithmically transformed), B. 2min digging latency in Vanilla DS+ mice from conventional (CH, n=7) or enriched (EH, n=9) housing (data logarithmically transformed), C. 1min digging latency in the same subjects (data Box Cox transformed).

Because only Vanilla mice met the requirement of treating the scent mixture as intermediate between the DS+ and DS-, simple effects of housing were calculated from the Trial Type*Housing term, using the SLICEDIFF command ([cf. 67]. Housing influenced digging latencies in the Vanilla DS+ mice, CH animals being slower than EH to dig in ambiguous trials (ambiguous: *t*=2.14, d.f.=91.89, *p*=0.018, *Cohen’s d*=1.083; positive: *t*=0.39, d.f.=91.89, *p*=0.348, *Cohen’s d*=0.198; negative: *t*=0.61, d.f.= 91.89, *p*=0.273, *Cohen’s d*=0.308: Fig. 3 B). Similar effect did not hold for Mint DS+ mice (ambiguous: *t*=0.68, d.f.=90.63, *p*=0.251, *Cohen’s d*=0.372; positive: *t*=-0.77, d.f.=90.63, *p*=0.221, *Cohen’s d*=-0.425; negative: *t*=0.75, d.f.= 90.63, *p*=0.229, *Cohen’s d*=0.410), even though Trial Type*Housing*DS+ odor and Trial Type*Housing did not account for significant variation (respectively *F*_2,65.37_=0.49, *p*=0.344 and *F*_2,62_=1.41, *p*=0.252). Digging duration was not affected by housing, in contrast (e.g. Vanilla DS+ ambiguous trials: *t*=-0.38, d.f.=91.37, *p*=0.353, *Cohen’s d*=-0.191).

Latency data were then re-analyzed using only the first minute of testing, to assess the utility of a shortened protocol. For Vanilla DS+ mice, SLICEDIFF tests for simple effects of Housing again showed that CH animals had longer latencies to dig in ambiguous trials than EH mice (ambiguous: *t*=2.27, d.f.=92.94, *p*=0.014, *Cohen’s d*=1.148; positive: *t*=0.22, d.f.= 92.94, *p*=0.414, *Cohen’s d*=0.110; negative: *t*=0.80, d.f.= 92.94, *p*=0.214, *Cohen’s d=0.404:* Fig. 3 C), while again the same did not hold for Mint DS+ mice (ambiguous: *t*=0.88, d.f.=91.94, *p*=0.193, *Cohen’s d*=0.482; positive: *t*=-0.65, d.f.= 91.94, *p*=0.260, *Cohen’s d*=-0.357; negative *t*=0.78, d.f.= 91.94, *p*=0.220, *Cohen’s d*=0.427). This was again despite Trial Type*Housing*DS+ odor and Trial Type*Housing not accounting for significant variation (respectively *F*2,65.37=0.36, *p*=0.392 and *F*_2,62_=1.66, *p*=0.198). Consistent with 2-minute results, C57s showed shorter latencies in the first minute (*F*_1,31_=8.49, *p*=0.003, *Cohen’s d*=-1.056) and there was a significant effect of Scented arm side (*F*_1,31_=6.81 *p*=0.001, *Cohen’s d*=-0.903).

Analyses of homecage observations confirmed expected housing effects on welfare (c.f. [38,69]): more stereotypic behavior (*F*_1,17.8_=25.19, *p*<0.0001, *Cohen’s d*=1.839) and time spent ‘inactive but awake’ (Wilcoxon rank sum test; *Z* =-2.839 *p*=0.008 *Cohen’s d*=0.484) in CH than EH cages. Taken together, these results validated digging latency as a JB indicator when vanilla is the DS+, and justified using a shortened, ‘ 1min’ protocol in Experiment 2.

### 2.4. Experiment 2: Applying the Task to Mice with Tumors

#### 2.4.1. Animals and Housing

Twenty male and 19 female ‘nude’ mice (stock NLAE:NIH(S)-*Fox1*^nu/nu^), free from common/zoonotic mouse pathogens, were obtained at 4-5 weeks old from La Plata National University’s Faculty of Veterinary Sciences and randomly allocated to cancer or cancer-free treatments (Fig. 4 B). We employed human A549 cell line lung adenocarcinoma tissue (from tumors grown in other mice: [70]): widely used in oncology [71] and associated with inflammatory cytokines [72]. The donor mouse was killed by cervical dislocation, and the tumor was aseptically removed and placed into a petri dish with Minimum Essential Medium, where it was divided into 2mm^2^ pieces. These pieces were immediately transplanted subcutaneously into the lateral abdominal area of mice anesthetized with ketamine/xylazine (100/10 mg/kg i.p., respectively). During the procedure, mice also received 0.03 ml of 1% lidocaine at the site of the incision and a single dose of 10 mg/kg of tramadol (administered subcutaneously). Post-operative care and ulterior veterinary inspection were performed as previously described [70].

**Fig 4:**
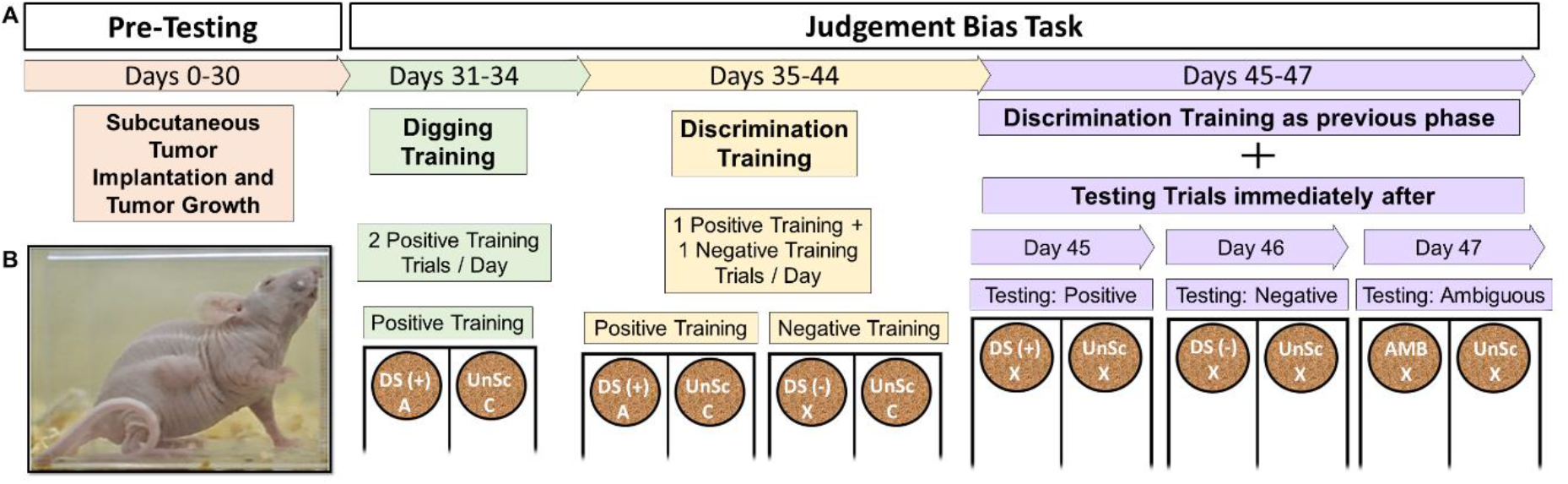
Timeline of positive, negative and ambiguous training and test trials for Experiment 2 (A), including a photograph of a mouse bearing a subcutaneous tumor (B). DS (+): positive discriminative stimulus (vanilla), DS (-): negative discriminative stimulus (mint), AMB: ambiguous mixture (50% vanilla-50%mint), A: almond, C: cornflake, X: no food rewards.

#### 2.4.2. Simplified JB protocol

Preference tests again identified high- and low-value rewards [73] (see Results). Training occurred on a clean bench in the colony room, under white light. A simplified protocol was used (Fig. 4 A, Table S2) in which vanilla was now used as the DS+ for all mice, since only vanilla DS+ mice in Experiment 1 interpreted intermediate mint-vanilla odor mixtures as ambiguous and responded to them with JB. Here, digging training was identical to Experiment 1 but lasted only 4 days, discrimination training lasted 9 days and involved 2, 3-minute trials per day, and testing lasted 3 days with mice meeting discrimination criteria being tested for responses to ambiguous cues as in Experiment 1. Daily veterinary checks assessed clinical signs of disease (c.f. [46])

#### 2.4.3. Statistical analyses for Experiment 2

Data were analyzed with repeated measures Generalized Linear Mixed Models as in Experiment 1, but model selection was not required due to the simpler design. Trial Type, Treatment (cancer status), Sex and their two- and 3-way interactions were therefore all included as fixed effects. Cage and Mouse ID, both nested within sex and treatment, were included as random effects. The simple effects of cancer status in each Trial Type were again tested using SLICEDIFF commands (now with two-tailed Ps). Sex differences in tumor volume and treat preference were also checked. For these, data did not meet assumptions of normality and homogeneity so a Wilcoxon rank sum test was used.

### 2.5. Experiment 2 Results

All but 3 male and 7 female mice met discrimination criteria (n=29). As for Experiment 1, high-value rewards were preferred over low-value rewards (males: Wilcoxon rank sum test; *Z* =-4.823, *p*<0.0001, *Cohen’s d*=0.234; females: Wilcoxon rank sum test; *Z* =-3.683, *p*=0.001, *Cohen’s d*=1.292). Trial Type*Treatment*Sex was significant (*F*_2,58.68_=8.77, *p*<0.001) because cancer status had different effects on males and females (Fig. 5 A, Table S4). In females, who treated the ambiguous cue as negative, only a trend for increased digging latency in negative trials for tumor-bearing animals was detected (*t*=-1.71, d.f.=80.65, *p*=0.093, *Cohen’s d*=1.001). In males, who in terms of latency treated the ambiguous cue as intermediate, the simple effect of cancer status was significant in ambiguous trials: tumor-bearing mice had longer latencies than controls (*t*=-2.93, d.f.=81.45, *p*=0.005, *Cohen’s d*=1.425). Unexpectedly, these mice also showed shorter latencies in negative trials (*t*=2.27, d.f.=81.83, *p*=0.027, *Cohen’s d*=-1.023). (Fig. 5 B).

**Fig 5.**
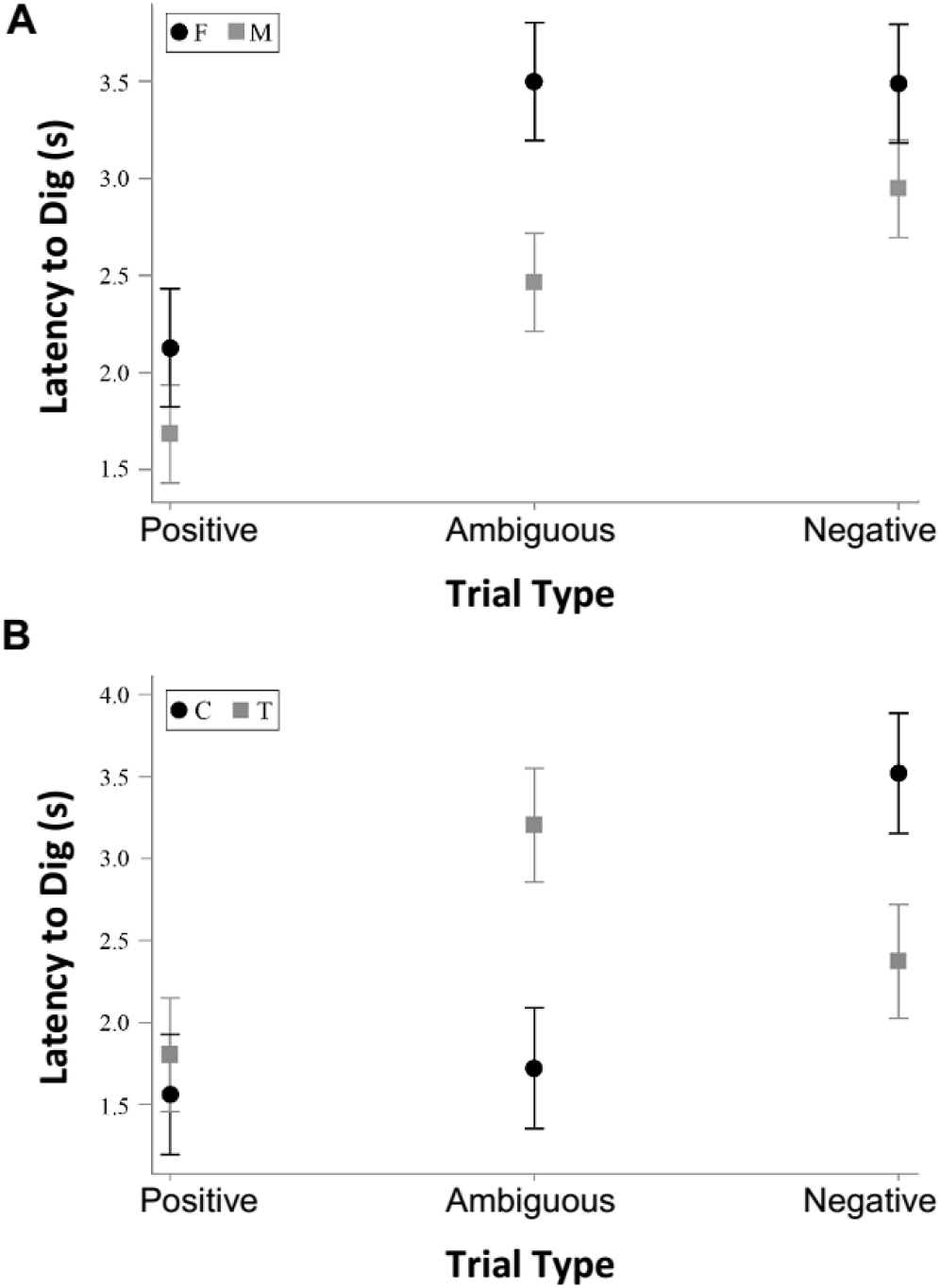
Digging latencies (± standard error) during positive, negative and ambiguous test trials (data logarithmically transformed). *A.* Digging latencies in female (F, n= 12) and male (M, n= 17) nude mice, *B.* Digging latencies in control (C, n=8) and tumor-bearing (T, n=9) males.

No mice presented clinical signs: all remained active with good body condition scores [74], normal gaits, normal skin condition over the tumors, and no nasal or ocular discharge. Following testing, tumor volumes were within accepted ranges [46], and similar between sexes (Wilcoxon rank sum test; *Z* =-0.7415, *p*=0.4698, *Cohen’s d*=1.292).

## 3. Discussion

To our knowledge, Experiment 1 represents the first successful construct validation of a mouse judgment bias (JB) task. Although sometimes omitted (5 studies in Table S1), or downplayed [26,75,76], the construct validation of any new indicator of affective state is a crucial step. This is especially important for animal JB tasks, which differ greatly from the unconditioned human tasks that inspired them [21] and which, perhaps as a consequence, can sometimes produce counterintuitive results [e.g. 77]. In mice, for instance, altered affective states have often failed to influence JB (10 studies in Table S1). We suspect that a combination of factors contributed to the success of this, the 11^th^ published validation attempt. First, affective state was manipulated through a highly preferred housing system that consistently produces robust differences in affect [34,36,38,39], resulting in large treatment effects. In these complex environments, with numerous places to hide or escape, stressful catching that otherwise could have masked treatment effects [78] was also avoided by training the mice to enter the ‘annex cages’. Second, the task was naturalistic and tailored to mouse olfactory and digging skills [79,80]. Third, we utilized a sensitive Go/Go design [21,22]; and rewards were pre-checked for their relative value. In addition, our initial counter-balanced design allowed us to detect an important asymmetry: when faced with a 50:50 mixture of scents, only mice trained to use vanilla as a DS+ treated this cue as ambiguous (a technical requirement for a valid JB task). Mice trained to use mint as the DS+ instead of treating these mixtures as 100% mint. This result identified Vanilla DS+ mice as the only potential candidates for validation of the current JB task (an issue we revisit below, in the context of future research directions).

For these Vanilla DS+ mice, latency to dig in ambiguous trials proved sensitive to well-being, thus showing construct validation: CH mice showed the predicted “pessimism”, taking longer to commence digging. Digging duration, however, proved insensitive to affect-modulated JB: although the expected graded response across trials was detected for this variable, no significant effect of housing was observed. This is perhaps because during ambiguous trials, subjects were influenced by the absence of reward, having learned in the positive and negative unreinforced test trials that if a treat was not detected early, further digging was not beneficial. However, it might also be that digging duration is only impacted by more severe affect manipulations (as we explore further below). Importantly, we were also able to demonstrate the versatility of this novel JB task across strains. Though strain differences were observed in Experiment 1, C57s being faster to begin digging while showing shorter digging durations, such effects were consistent across trial type, and housing treatments. This suggests that when technical criteria are met -- mice being able to discriminate between positive and negative cues, and interpreting intermediate cues as ambiguous -- the task has potential generalizability across strains. Moreover, the nude mice of Experiment 2 (with a Swiss genetic background), were also able to meet these technical criteria even after a shortened protocol.

In Experiment 2, an abbreviated form of this task (the labor-intensiveness of JB testing being a challenge [81,82]) thus tested hypotheses about the affective impacts of cancer. In males faced with ambiguous cues, tumor implantation reduced digging latencies indicating negative JBs. This, the first evidence of pessimism in tumor-bearing animals, is consistent with the low mood commonly reported in human patients and the depression-like behavioral changes commonly found in rodent models (see Introduction). The degree of pessimism in tumor-bearing males was surprising however, ambiguous cues were treated even more negatively than negative trials. This effect may parallel the influence that human affective states have on response latency during JB tasks: individuals in some negative states are slower to make decisions when presented with ambiguous cues [83,84]. It is also unclear why females showed no tumor-induced JB. It is possible that this reflects genuine sex differences (since male mice seem more adversely affected than females later in disease progression [85]). But this might instead indicate floor effects, since even control females seemed to treat ambiguous cues as negative, perhaps because all female mice were experiencing negative affect from being kept in barren CH (c.f. Experiment 1). The results from tumor-bearing mice thus confirmed our new task’s potential for assessing affective state in biomedical research. But they also highlighted needs for further validation and refinement.

Together, our results thus identify a valid JB task with great promise, but also show that its sensitivity needs improving. Future avenues could include reducing potential floor effects in replicate cancer research by housing nude mice with enrichment [c.f. 73]. Reinvestigating the construct validity of digging duration would also be warranted. Incidental findings revealed that despite Experiment 1’s null results, Experiment 2’s tumor-bearing males (pessimistic according to JB latencies) also showed shorter digging durations in ambiguous trials, suggesting that this measure too was sensitive to low mood (see Figure S1 and Table S5). Future studies could thus evaluate more negative modifications of affective state in replicate validation work, to assess whether digging duration is sensitive to these more challenging manipulations (Experiment 1’s null results then being floor effects). Experimenting with different training odors, and also different mixtures as ambiguous cues, is also now warranted. To illustrate, Experiment 1’s Mint DS+ mice treated 50/50 mixtures as 100% mint (i.e. positive) and even to human noses these mixtures smelled strongly ‘minty’. This suggests that when using a mint DS+, more vanilla-skewed mixtures are needed for them to be treated as intermediate. Further, when faced with the same mixtures, female nude mice trained with Vanilla as DS+ treated them as *negative,* suggesting that for these mice, ‘near positive’ ambiguous cues skewed toward vanilla would be needed for a mixture to be treated as intermediate. Pilot tests to identify appropriate odor mixtures, and/or use of multiple ambiguous probes could help mitigate such effects. Exposing subjects to a full spectrum of ambiguous cues (e.g. near positive, intermediate, near negative) could also allow for distinction between different types of negative states (e.g. anxiety- and depression-like responses, see [18,21]). And as a final avenue for future research, since housing and cancer status are both long term manipulations, future work assessing the sensitivity of this JB task to acute changes in affect would be of great benefit.

Such methodological refinements are important given the great research value of a validated murine JB task. As a humane technique potentially sensitive to both positive and negative affect [11,86], it could be hugely useful for assessing and improving mouse husbandry. Further, cognitive biases play important roles in human affective disorders, yet relatively little is known about the neurophysiological correlates and underlying mechanisms [84]. Since mice are the dominant model species in cognitive and behavioral neuroscience, a validated murine JB task opens the doors to further investigation and understanding of these mechanisms. It could be valuable in biomedical research too. In oncology, for instance, a validated JB task could help investigate how cancer influences affective states (as in our preliminary work). But it could also help investigate the contradictory effects of opiates on cancer pathogenicity [c.f. 87]; the adverse effects of chemotherapy on wellbeing [88]; and the role of mood in the cancer-protective effects of housing rodents with warmth [89], companions [90] and enrichments [91]: fascinating topics with both animal welfare and clinical implications.

## 4. Conclusion

In summary, this novel JB task has proven to be a valid indicator of affective state. For C57 and Balb females, CH animals showed predicted pessimistic responses to ambiguous cues when their latency to dig was assessed. For tumor-bearing male nude mice, this task also indicated negative states through pessimistic responses to ambiguity, even before the occurrence of clinical signs of disease. Together, these results highlight the potential value of this novel murine JB task across diverse fields of research. This indicator of affective state is easy to implement, economic, and has already proven effective in three common mouse strains. But in addition to its utility, this JB task is also humane: an uncommon feature in available indicators of mouse affect, which often involve aversive experiences (e.g. exposure to open fields, elevated platforms, electrical shocks). This JB task thus offers a welfare-friendly alternative to standard indicators of affect. Although replication and further refinements to improve sensitivity are still needed, a valid JB task has potential in animal welfare assessment and addressing fundamental questions about affective state.

## 5. Acknowledgments

The authors would like to acknowledge that Experiment 1 was completed on the ancestral lands of the Attawandaron people and the treaty lands and territory of the Mississaugas of the Credit. The authors would also like to thank the mice, Dr. Jim Petrick for providing comments on the manuscript and our wonderful animal care technicians Michaela Randal and Michelle Cieplak.

## References

[1] J. Keen, Wasted money in United States biomedical and agricultural animal research, in: K. Herrmann, K. Jayne (Eds.), Animal Experimentation: Working Towards a Paradigm Change, Brill, Leiden, The Netherlands, 2019: pp. 244–272.

[2] K. Gouveia, J.L. Hurst, Optimising reliability of mouse performance in behavioural testing: The major role of non-aversive handling, Sci. Rep. 7 (2017) 44999. https://doi.org/10.1038/srep44999.

[3] J.P. Balcombe, Laboratory environments and rodents’ behavioural needs: A review, Lab. Anim. 40 (2006) 217–235. https://doi.org/10.1258/002367706777611488.

[4] V. Baumans, The impact of the environment on laboratory animals, in: M.L. Andersen, S. Tufik (Eds.), Rodent Model as Tools in Ethical Biomedical Research, Springer, 2016: pp. 13–22.

[5] I.A.S. Olsson, K. Dahlborn, Improving housing conditions for laboratory mice: A review of “environmental enrichment,” Lab. Anim. 36 (2002) 243–270. https://doi.org/10.1258/002367702320162379.

[6] T. Poole, Happy animals make good science, Lab. Anim. 31 (1997) 116–124. https://doi.org/10.1258/002367797780600198.

[7] L.P. Freedman, I.M. Cockburn, T.S. Simcoe, The economics of reproducibility in preclinical research, PLoS Biol. 13 (2015) e1002165. https://doi.org/10.1371/journal.pbio.1002165.

[8] I.W.Y. Mak, N. Evaniew, M. Ghert, Lost in translation: Animal models and clinical trials in cancer treatment, Am. J. Transl. Res. 6 (2014) 114–118.

[9] C.H. Wong, K.W. Siah, A.W. Lo, Estimation of clinical trial success rates and related parameters, Biostatistics. 20 (2019) 273–286. https://doi.org/10.1093/biostatistics/kxx069.

[10] J.P. Garner, The significance of meaning: Why do over 90% of behavioral neuroscience results fail to translate to humans, and what can we do to fix it?, ILAR J. 55 (2014) 438–456. https://doi.org/10.1093/ilar/ilu047.

[11] M. Mendl, O.H.P. Burman, R.M.A. Parker, E.S. Paul, Cognitive bias as an indicator of animal emotion and welfare: Emerging evidence and underlying mechanisms, Appl. Anim. Behav. Sci. 118 (2009) 161–181. https://doi.org/10.1016/j.applanim.2009.02.023.

[12] European Commission, Report From the Commission to the European Parlaiment and the Council, 2020. https://eur-lex.europa.eu/legal-content/EN/TXT/HTML/?uri=CELEX:52020DC0016&from=EN.

[13] I. Blanchette, A. Richards, The influence of affect on higher level cognition: A review of research on interpretation, judgement, decision making and reasoning, Cogn. Emot. 24 (2010) 561–595. https://doi.org/10.1080/02699930903132496.

[14] C. MacLeod, I.L. Cohen, Anxiety and the Interpretation of Ambiguity: A Text Comprehension Study, J. Abnorm. Psychol. 102 (1993) 238–247. https://doi.org/10.1037/0021-843X.102.2.238.

[15] A. Mathews, C. MacLeod, Cognitive approaches to emotions, Anu. Rev. Psychol. 45 (1994) 25–50.

[16] C. Douglas, M. Bateson, C. Walsh, A. Bédué, S.A. Edwards, Environmental enrichment induces optimistic cognitive biases in pigs, Appl. Anim. Behav. Sci. 139 (2012) 65–73. https://doi.org/10.1016/j.applanim.2012.02.018.

[17] E.J. Harding, E.S. Paul, M. Mendl, Cognitive bias and affective state, Nature. 427 (2004) 312. https://doi.org/10.1038/427312a.

[18] M. Mendl, O.H.P. Burman, R.M.A. Parker, E.S. Paul, Cognitive bias as an indicator of animal emotion and welfare: Emerging evidence and underlying mechanisms, Appl. Anim. Behav. Sci. 118 (2009) 161–181. https://doi.org/10.1016/j.applanim.2009.02.023.

[19] M. Mendl, E. Paul, Getting to the heart of animal welfare. The study of animal emotion, Stichting Animales, Netherlands, 2017.

[20] L. Gygax, The A to Z of statistics for testing cognitive judgement bias, Anim. Behav. 95 (2014) 59–69. https://doi.org/10.1016/j.anbehav.2014.06.013.

[21] S. Roelofs, H. Boleij, R.E. Nordquist, F.J. van der Staay, Making decisions under ambiguity: Judgment bias tasks for assessing emotional state in animals, Front. Behav. Neurosci. 10 (2016) 1–16. https://doi.org/10.3389/fnbeh.2016.00119.

[22] M. Lagisz, J. Zidar, S. Nakagawa, V. Neville, E. Sorato, E.S. Paul, M. Bateson, M. Mendl, H. Løvlie, Optimism, pessimism and judgement bias in animals: A systematic review and meta-analysis, Neurosci. Biobehav. Rev. 118 (2020) 3–17. https://doi.org/10.1016/j.neubiorev.2020.07.012.

[23] J. Ahloy-Dallaire, J. Espinosa, G. Mason, Play and optimal welfare: Does play indicate the presence of positive affective states?, Behav. Processes. 156 (2018) 3–15. https://doi.org/10.1016/j.beproc.2017.11.011.

[24] S. Jones, E.S. Paul, P. Dayan, E.S.J. Robinson, M. Mendl, Pavlovian influences on learning differ between rats and mice in a counter-balanced Go/NoGo judgement bias task, Behav. Brain Res. 331 (2017) 214–224. https://doi.org/10.1016/j.bbr.2017.05.044.

[25] S. Hintze, L. Melotti, S. Colosio, J.D. Bailoo, M. Boada-Saña, H. Würbel, E. Murphy, A cross-species judgement bias task: Integrating active trial initiation into a spatial Go/No-go task, Sci. Rep. 8 (2018) 1–13. https://doi.org/10.1038/s41598-018-23459-3.

[26] V. Krakenberg, I. Woigk, L. Garcia Rodriguez, N. Kästner, S. Kaiser, N. Sachser, S.H. Richter, Technology or ecology? New tools to assess cognitive judgement bias in mice, Behav. Brain Res. 362 (2019) 279–287. https://doi.org/10.1016/j.bbr.2019.01.021.

[27] H. Boleij, J. van’t Klooster, M. Lavrijsen, S. Kirchhoff, S.S. Arndt, F. Ohl, A test to identify judgement bias in mice, Behav. Brain Res. 233 (2012) 45–54. https://doi.org/10.1016/j.bbr.2012.04.039.

[28] V. Kloke, R.S. Schreiber, C. Bodden, J. Möllers, H. Ruhmann, S. Kaiser, K.P. Lesch, N. Sachser, L. Lewejohann, Hope for the best or prepare for the worst? Towards a spatial cognitive bias test for mice, PLoS One. 9 (2014) e105431. https://doi.org/10.1371/journal.pone.0105431.

[29] J. Novak, J.D. Bailoo, L. Melotti, J. Rommen, H. Würbel, An exploration based cognitive bias test for mice: Effects of handling method and stereotypic behaviour, PLoS One. 10 (2015) e0130718. https://doi.org/10.1371/journal.pone.0130718.

[30] J. Novak, K. Stojanovski, L. Melotti, T. Reichlin, R. Palme, H. Würbel, Effects of stereotypic behaviour and chronic mild stress on judgement bias in laboratory mice, Appl. Anim. Behav. Sci. 174 (2016) 162–172. https://doi.org/10.1016/j.applanim.2015.10.004.

[31] J. Novak, J.D. Bailoo, L. Melotti, H. Würbel, Effect of cage-induced stereotypies on measures of affective state and recurrent perseveration in CD-1 and C57BL/6 mice, PLoS One. 11 (2016) e0153203. https://doi.org/10.1371/journal.pone.0153203.

[32] J.D. Bailoo, E. Murphy, M. Boada-Saña, J.A. Varholick, S. Hintze, C. Baussière, K.C. Hahn, C. Göpfert, R. Palme, B. Voelkl, H. Würbel, Effects of cage enrichment on behavior, welfare and outcome variability in female mice, Front. Behav. Neurosci. 12 (2018). https://doi.org/10.3389/fnbeh.2018.00232.

[33] V. Krakenberg, V.T. von Kortzfleisch, S. Kaiser, N. Sachser, S.H. Richter, Differential Effects of Serotonin Transporter Genotype on Anxiety-Like Behavior and Cognitive Judgment Bias in Mice, Front. Behav. Neurosci. 13 (2019) 263. https://doi.org/10.3389/fnbeh.2019.00263.

[34] S.C. Tilly, J. Dallaire, G.J. Mason, Middle-aged mice with enrichment-resistant stereotypic behaviour show reduced motivation for enrichment, Anim. Behav. 80 (2010) 363–373. https://doi.org/10.1016/j.anbehav.2010.06.008.

[35] S. Chourbaji, C. Zacher, C. Sanchis-Segura, R. Spanagel, P. Gass, Social and structural housing conditions influence the development of a depressive-like phenotype in the learned helplessness paradigm in male mice, Behav. Brain Res. 164 (2005) 100–106. https://doi.org/10.1016/j.bbr.2005.06.003.

[36] A. Adcock, E. Choleris, M. Denommé, H. Khan, L. Levison, A. MacLellan, B. Nazal, L. Niel, E. Nip, G. Mason, Where are you from? Female mice raised in enriched or conventional cages differ socially, and can be discriminated by other mice, Behav. Brain Res. 400 (2021) 113025. https://doi.org/10.1016/j.bbr.2020.113025.

[37] N. Benaroya-Milshtein, N. Hollander, A. Apter, T. Kukulansky, N. Raz, A. Wilf, I. Yaniv, C.G. Pick, Environmental enrichment in mice decreases anxiety, attenuates stress responses and enhances natural killer cell activity, Eur. J. Neurosci. 20 (2004) 1341–1347. https://doi.org/10.1111/j.1460-9568.2004.03587.x.

[38] C. Fureix, M. Walker, L. Harper, K. Reynolds, A. Saldivia-woo, G. Mason, Stereotypic behaviour in standard non-enriched cages is an alternative to depression-like responses in C57BL/6 mice, Behav. Brain Res. 305 (2016) 186–190. https://doi.org/10.1016/j.bbr.2016.02.005.

[39] E. Nip, A. Adcock, B. Nazal, A. MacLellan, L. Niel, E. Choleris, L. Levison, G. Mason, Why are enriched mice nice ? Investigating how environmental enrichment reduces agonism in female C57BL / 6, DBA / 2, and BALB / c mice, Appl. Anim. Behav. Sci. 217 (2019) 73–82. https://doi.org/10.1016/j.applanim.2019.05.002.

[40] V. Baumans, The laboratory mouse, in: R.C. Hubrecht, J. Kirkwood (Eds.), UFAW Handbook on the Care and Management of Laboratory and Other Research Animals, 8th ed., Wiley-Blackwell, Oxford, UK, 2010: pp. 276–310.

[41] G. Kempermann, Environmental enrichment, new neurons and the neurobiology of individuality, Nat. Rev. Neurosci. 20 (2019) 235–245. https://doi.org/10.1038/s41583-019-0120-x.

[42] C. Poirier, M. Bateson, F. Gualtieri, E.A. Armstrong, G.C. Laws, T. Boswell, T. V. Smulders, Validation of hippocampal biomarkers of cumulative affective experience, Neurosci. Biobehav. Rev. 101 (2019) 113–121. https://doi.org/10.1016/j.neubiorev.2019.03.024.

[43] S.H. Richter, A. Schick, C. Hoyer, K. Lankisch, P. Gass, B. Vollmayr, A glass full of optimism: Enrichment effects on cognitive bias in a rat model of depression, Cogn. Affect. Behav. Neurosci. 12 (2012) 527–542. https://doi.org/10.3758/s13415-012-0101-2.

[44] N.M. Brydges, M. Leach, K. Nicol, R. Wright, M. Bateson, Environmental enrichment induces optimistic cognitive bias in rats, Anim. Behav. 81 (2011) 169–175. https://doi.org/10.1016/j.anbehav.2010.09.030.

[45] J.V. Roughan, C.A. Coulter, P.A. Flecknell, H.D. Thomas, K.J. Sufka, The conditioned place preference test for assessing welfare consequences and potential refinements in a mouse bladder cancer model, PLoS One. 9 (2014) e103362. https://doi.org/10.1371/journal.pone.0103362.

[46] P. Workman, E.O. Aboagye, F. Balkwill, A. Balmain, G. Bruder, D.J. Chaplin, J.A. Double, J. Everitt, D.A.H. Farningham, M.J. Glennie, L.R. Kelland, V. Robinson, I.J. Stratford, G.M. Tozer, S. Watson, S.R. Wedge, S.A. Eccles, V. Navaratnam, S. Ryder, Guidelines for the welfare and use of animals in cancer research, Br. J. Cancer. 102 (2010) 1555–1577. https://doi.org/10.1038/sj.bjc.6605642.

[47] A. Schrepf, S.K. Lutgendorf, L.M. Pyter, Pre-treatment effects of peripheral tumors on brain and behavior: Neuroinflammatory mechanisms in humans and rodents, Brain. Behav. Immun. 49 (2015) 1–17. https://doi.org/10.1016/j.bbi.2015.04.010.

[48] R.J. Dunlop, C.W. Campbell, Cytokines and advanced cancer, J. Pain Symptom Manage. 20 (2000) 214–232. https://doi.org/10.1016/S0885-3924(00)00199-8.

[49] L. Van Esch, J.A. Roukema, M.F. Ernst, G.A.P. Nieuwenhuijzen, J. De Vries, Combined anxiety and depressive symptoms before diagnosis of breast cancer, J. Affect. Disord. 136 (2012) 895–901. https://doi.org/10.1016/j.jad.2011.09.012.

[50] G.D. Friedman, J.S. Skilling, N. V. Udaltsova, L.H. Smith, Early symptoms of ovarian cancer: A case-control study without recall bias, Fam. Pract. 22 (2005) 548–553. https://doi.org/10.1093/fampra/cmi044.

[51] B. Bortolato, T.N. Hyphantis, S. Valpione, G. Perini, M. Maes, G. Morris, M. Kubera, C.A. Köhler, B.S. Fernandes, B. Stubbs, N. Pavlidis, A.F. Carvalho, Depression in cancer: The many biobehavioral pathways driving tumor progression, Cancer Treat. Rev. 52 (2017) 58–70. https://doi.org/10.1016/j.ctrv.2016.11.004.

[52] F. Colotta, P. Allavena, A. Sica, C. Garlanda, A. Mantovani, Cancer-related inflammation, the seventh hallmark of cancer: Links to genetic instability, Carcinogenesis. 30 (2009) 1073–1081. https://doi.org/10.1093/carcin/bgp127.

[53] M.G. Nashed, E.P. Seidlitz, B.N. Frey, G. Singh, Depressive-like behaviours and decreased dendritic branching in the medial prefrontal cortex of mice with tumors: A novel validated model of cancer-induced depression, Behav. Brain Res. 294 (2015) 25–35. https://doi.org/10.1016/j.bbr.2015.07.040.

[54] D.M. Norden, R. Devine, S. Bicer, R. Jing, P.J. Reiser, L.E. Wold, J.P. Godbout, D.O. McCarthy, Fluoxetine prevents the development of depressive-like behavior in a mouse model of cancer related fatigue, Physiol. Behav. 140 (2015) 230–235. https://doi.org/10.1016/j.physbeh.2014.12.045.

[55] D.M. Lamkin, S.K. Lutgendorf, D. Lubaroff, A.K. Sood, T.G. Beltz, A.K. Johnson, Cancer induces inflammation and depressive-like behavior in the mouse: Modulation by social housing, Brain. Behav. Immun. 25 (2011) 555–564. https://doi.org/10.1016/j.bbi.2010.12.010.

[56] M. Yang, J. Kim, J.S. Kim, S.H. Kim, J.C. Kim, M.J. Kang, U. Jung, T. Shin, H. Wang, C. Moon, Hippocampal dysfunctions in tumor-bearing mice, Brain. Behav. Immun. 36 (2014) 147–155. https://doi.org/10.1016/j.bbi.2013.10.022.

[57] A.M. Casaril, M. Domingues, S.R. Bampi, D. de Andrade Lourenço, T.Â. Smaniotto, N. Segatto, B. Vieira, F.K. Seixas, T. Collares, E.J. Lenardão, L. Savegnago, The antioxidant and immunomodulatory compound 3-[(4-chlorophenyl)selanyl]-1-methyl-1H-indole attenuates depression-like behavior and cognitive impairment developed in a mouse model of breast tumor, Brain. Behav. Immun. 84 (2020) 229–241. https://doi.org/10.1016/j.bbi.2019.12.005.

[58] D.M. Norden, S. Bicer, Y. Clark, R. Jing, C.J. Henry, L.E. Wold, P.J. Reiser, J.P. Godbout, D.O. McCarthy, Tumor growth increases neuroinflammation, fatigue and depressive-like behavior prior to alterations in muscle function, Brain. Behav. Immun. 43 (2015) 76–85. https://doi.org/10.1016/j.bbi.2014.07.013.

[59] N. Percie du Sert, V. Hurst, A. Ahluwalia, S. Alam, M.T. Avey, M. Baker, W.J. Browne, A. Clark, I.C. Cuthill, U. Dirnagl, M. Emerson, P. Garner, S.T. Holgate, D.W. Howells, N.A. Karp, S.E. Lazic, K. Lidster, C.J. MacCallum, M. Macleod, E.J. Pearl, O.H. Petersen, F. Rawle, P. Reynolds, K. Rooney, E.S. Sena, S.D. Silberberg, T. Steckler, H. Würbel, The ARRIVE guidelines 2.0: Updated guidelines for reporting animal research*, J. Cereb. Blood Flow Metab. 40 (2020) 1769–1777. https://doi.org/10.1177/0271678X20943823.

[60] E.M. Weber, J.A. Dallaire, B.N. Gaskill, K.R. Pritchett-Corning, J.P. Garner, Aggression in group-housed laboratory mice: Why can’t we solve the problem?, Lab Anim. (NY). 46 (2017) 157–161. https://doi.org/10.1038/laban.1219.

[61] M. Walker, C. Fureix, R. Palme, J.A. Newman, J. Ahloy Dallaire, G. Mason, Mixed-strain housing for female C57BL/6, DBA/2, and BALB/c mice: Validating a split-plot design that promotes refinement and reduction, BMC Med. Res. Methodol. 16 (2016). https://doi.org/10.1186/s12874-016-0113-7.

[62] M. Haahr, RANDOM.ORG: True Random Number Service, (2021). https://www.random.org.

[63] J. Chapuis, D.A. Wilson, Cholinergic modulation of olfactory pattern separation, Neurosci. Lett. 545 (2013) 50–53. https://doi.org/10.1016/j.neulet.2013.04.015.

[64] J. Chapuis, Y. Cohen, X. He, Z. Zhang, S. Jin, F. Xu, D.A. Wilson, Lateral entorhinal modulation of piriform cortical activity and fine odor discrimination, J. Neurosci. 33 (2013) 13449–13459. https://doi.org/10.1523/JNEUROSCI.1387-13.2013.

[65] L. Harper, E. Choleris, K. Ervin, C. Fureix, K. Reynolds, M. Walker, G. Mason, Stereotypic mice are aggressed by their cagemates, and tend to be poor demonstrators in social learning tasks, Anim. Welf. 24 (2015) 463–473.

[66] K.P. Burnham, D.R. Anderson, Model selection and multimodel inference: a practical information-theoretic approach, 2nd ed., Springer, New York, 2002.

[67] J. Wei, R.J. Carroll, K.K. Harden, G. Wu, Comparisons of treatment means when factors do not interact in two-factorial studies, Amino Acids. 42 (2012) 2031–2035. https://doi.org/10.1007/s00726-011-0924-0.

[68] G.D. Ruxton, M. Neuhäuser, When should we use one-tailed hypothesis testing?, Methods Ecol. Evol. 1 (2010) 114–117. https://doi.org/10.1111/j.2041-210x.2010.00014.x.

[69] G. Mason, J. Rushen, eds., Stereotypic Animal Behaviour—Fundamentals and Applications to Welfare, 2nd ed., CABI, Wallingford, Oxford, 2006.

[70] A. Resasco, A.C. Carranza Martin, M.A. Ayala, S.L. Diaz, C. Carbone, Non-aversive photographic measurement method for subcutaneous tumours in nude mice, Lab. Anim. 53 (2019) 352–361. https://doi.org/10.1177/0023677218793450.

[71] L. Korrodi-Gregório, V. Soto-Cerrato, R. Vitorino, M. Fardilha, R. Pérez-Tomás, From proteomic analysis to potential therapeutic targets: Functional profile of two lung cancer cell lines, A549 and SW900, widely studied in pre-clinical research, PLoS One. 11 (2016) e0165973. https://doi.org/10.1371/journal.pone.0165973.

[72] Q. Huang, L. Duan, X. Qian, J. Fan, Z. Lv, X. Zhang, J. Han, F. Wu, M. Guo, G. Hu, J. Du, C. Chen, Y. Jin, IL-17 Promotes Angiogenic Factors IL-6, IL-8, and Vegf Production via Stat1 in Lung Adenocarcinoma, Sci. Rep. 6 (2016) 36551. https://doi.org/10.1038/srep36551.

[73] A. Resasco, Impacto del desarrollo de la línea tumoral A549 en el bienestar de ratones de la cepa NLAE:NIH(S)-*Fox1*nu/nu, Universidad Nacional De La Plata, 2018. http://sedici.unlp.edu.ar/bitstream/handle/10915/87590/Documento_completo.pdf?sequence=1.

[74] M.H. Ullman-Culleré, C.J. Foltz, Body condition scoring: A rapid and accurate method for assessing health status in mice, Lab. Anim. Sci. 49 (1999) 319–323.

[75] A. Verjat, P. Devienne, H.G. Rödel, C. Féron, More exploratory house mice judge an ambiguous situation more negatively, Anim. Cogn. 24 (2021) 53–64. https://doi.org/10.1007/s10071-020-01414-y.

[76] H.A.T. Nguyen, C. Guo, J.R. Homberg, Cognitive Bias Under Adverse and Rewarding Conditions: A Systematic Review of Rodent Studies, Front. Behav. Neurosci. 14 (2020) 1–12. https://doi.org/10.3389/fnbeh.2020.00014.

[77] E. Verbeek, D. Ferguson, C. Lee, Are hungry sheep more pessimistic? The effects of food restriction on cognitive bias and the involvement of ghrelin in its regulation, Physiol. Behav. 123 (2014) 67–75. https://doi.org/10.1016/j.physbeh.2013.09.017.

[78] E. Nip, The Long-Term Effects of Environmental Enrichment on Agonism in Female C57BL/6, BALB/c, and DBA/2 Mice, University of Guelph, 2018.

[79] N. Latham, G. Mason, From house mouse to mouse house: The behavioural biology of free-living Mus musculus and its implications in the laboratory, Appl. Anim. Behav. Sci. 86 (2004) 261–289. https://doi.org/10.1016/j.applanim.2004.02.006.

[80] J.W. Young, L.E. Kerr, J.S. Kelly, H.M. Marston, C. Spratt, K. Finlayson, J. Sharkey, The odour span task: A novel paradigm for assessing working memory in mice, Neuropharmacology. 52 (2007) 634–645. https://doi.org/10.1016/j.neuropharm.2006.09.006.

[81] S. Jones, V. Neville, L. Higgs, E.S. Paul, P. Dayan, E.S.J. Robinson, M. Mendl, Assessing animal affect: an automated and self-initiated judgement bias task based on natural investigative behaviour, Sci. Rep. 8 (2018) 12400. https://doi.org/10.1038/s41598-018-30571-x.

[82] N.M. Brydges, L. Hall, A shortened protocol for assessing cognitive bias in rats, J. Neurosci. Methods. 286 (2017) 1–5. https://doi.org/10.1016/j.jneumeth.2017.05.015.

[83] E.S. Paul, I. Cuthill, G. Kuroso, V. Norton, J. Woodgate, M. Mendl, Mood and the speed of decisions about anticipated resources and hazards, Evol. Hum. Behav. 32 (2011) 21–28. https://doi.org/10.1016/j.evolhumbehav.2010.07.005.

[84] A. Schick, M. Wessa, B. Vollmayr, C. Kuehner, P. Kanske, Indirect assessment of an interpretation bias in humans: Neurophysiological and behavioral correlates, Front. Hum. Neurosci. 7 (2013) 1–11. https://doi.org/10.3389/fnhum.2013.00272.

[85] P.F. Cosper, L.A. Leinwand, Cancer causes cardiac atrophy and autophagy in a sexually dimorphic manner, Cancer Res. 71 (2011) 1710–1720. https://doi.org/10.1158/0008-5472.CAN-10-3145.

[86] E.S. Paul, E.J. Harding, M. Mendl, Measuring emotional processes in animals: The utility of a cognitive approach, Neurosci. Biobehav. Rev. 29 (2005) 469–491. https://doi.org/10.1016/j.neubiorev.2005.01.002.

[87] B. Afsharimani, K. Kindl, P. Good, J. Hardy, Pharmacological options for the management of refractory cancer pain—what is the evidence?, Support. Care Cancer. 23 (2015) 1473–1481. https://doi.org/10.1007/s00520-015-2678-9.

[88] L.J. Wood, L.M. Nail, A. Gilster, K.A. Winters, C.R. Elsea, Cancer chemotherapy-related symptoms: Evidence to suggest a role for proinflammatory cytokines, Oncol. Nurs. Forum. 33 (2006) 536–542. https://doi.org/10.1188/06.ONF.535-542.

[89] K.M. Kokolus, M.L. Capitano, C.T. Lee, J.W.L. Eng, J.D. Waight, B.L. Hylander, S. Sexton, C.C. Hong, C.J. Gordon, S.I. Abrams, E.A. Repasky, Baseline tumor growth and immune control in laboratory mice are significantly influenced by subthermoneutral housing temperature, Proc. Natl. Acad. Sci. U. S. A. 110 (2013) 20176–20181. https://doi.org/10.1073/pnas.1304291110.

[90] G.L. Hermes, B. Delgado, M. Tretiakova, S.A. Cavigelli, T. Krausz, S.D. Conzen, M.K. McClintock, Social isolation dysregulates endocrine and behavioral stress while increasing malignant burden of spontaneous mammary tumors, Proc. Natl. Acad. Sci. U. S. A. 106 (2009) 22393–22398. https://doi.org/10.1073/pnas.0910753106.

[91] L. Cao, X. Liu, E.J.D. Lin, C. Wang, E.Y. Choi, V. Riban, B. Lin, M.J. During, Environmental and Genetic Activation of a Brain-Adipocyte BDNF/Leptin Axis Causes Cancer Remission and Inhibition, Cell. 142 (2010) 52–64. https://doi.org/10.1016/j.cell.2010.05.029.

