## Supplementary material for "Cancer blues? A validated judgment bias task suggests pessimism in nude mice with tumors"

**Table S1:** Summary of judgement bias assays for mice, outlining task design, requirements for construct validation and whether task was validated as an indicator of affective state

| Study | Test Type | Subjects | Discriminative stimuli (DS) | DS+ / DS- counterbalanced? | Unconditioned stimuli (UCS) | Trained response | Discrimination criterion (DC) | Successful Discrimination? | Affective state manipulation / inference used in construct validation | Justification / confirmation of affective state manipulation | Were intermediate cues perceived as ambiguous? | Treatment effects on response(s) to ambiguous cues | Required response to ambiguity for successful construct validation | Successful construct validation? (i.e. test is sensitive to affective state) | Reason for failed validation |
|---|---|---|---|---|---|---|---|---|---|---|---|---|---|---|---|
| **Boleij et al., (2012) Exp 1** | Olfactory (Go/No-Go) | 50 male Balb/cJ<br><br>50 male 129P3J | DS+/DS-: Vanilla/apple essence (0.05%)<br><br>AMB: 85%POS15%NEG (near positive)<br><br>50%POS50%NEG (ambiguous)<br><br>15%POS85%NEG (near negative) | Yes | UCS+: Palatable Almond<br><br>UCS-: Unpalatable Almond | Eat palatable almond (POS) or not eat unpalatable almond (NEG) | Group level only: statistically significant differences between POS and NEG | Yes (only Balb/cs, not 129P3s) | Strain differences: high (Balb/cJ) or low (129P3J) trait anxiety | Well documented increased anxiety-like behavior in Balb/c mice in literature | Yes (only Balb/cs, not 129P3s) | No strain difference: 129P3 mice could not discriminate the task | Longer latency to eat the almond for Balb/c mice | No | 129P3 mice did not discriminate between type of trial |
| **Boleij et al., (2012) Exp 2** | Olfactory (Go/No-Go) | 50 male Balb/cJ | DS+/DS-: Vanilla/apple essence (0.05%)<br><br>AMB: 50%POS50%NEG | Yes | UCS+: Palatable Almond<br><br>UCS-: Unpalatable Almond | Eat palatable almond (POS) or not eat unpalatable almond (NEG) | Group level only: statistically significant differences between POS and NEG | Yes | Testing under bright (aversive) vs. dark (non-aversive) lighting conditions | Well documented mouse aversion to white light in literature | No: AMB cue was perceived as NEG for both light conditions | Light treatment effect not specific to AMB trial: increase in latency in mice tested under white light for all trials | Longer latency to eat the almond under white light | No | White light increased latencies for all trials, and ambiguous cue was interpreted as negative for all mice. |
| **Kloke et al., (2014) Exp 3** | Spatial (Go/No-Go) | 12 female 5HTT (+/+)<br><br>12 female 5HTT (+/-)<br><br>12 female 5HTT (-/-) | DS+/DS-: Location in a three-arm radial maze<br><br>AMB: Middle arm between POS and NEG | Yes | UCS+: Homecage access<br><br>UCS-: Air-puff | Enter (POS) or not enter (NEG) the hole at the end of a radial maze arm | Shorter latencies in all positive training trials compared to all negative training trials on day 4 of training | Yes (55% of mice) | Compared serotonin transporter (5-HTT) knockout mice: wild-type 5HTT (+/+)<br><br>heterozygous 5HTT (+/-)<br><br>homozygous 5HTT (-/-) | Increased anxiety- and depression-like behaviors in 5-HTT knockout mice documented in literature | N/A: AMB arm not directly compared to POS and NEG | Non-significant trend for strain effect | Longer AMB latency in homozygous 5-HTT knockout mice than heterozygous, followed by wild-type mice | No | Differences between strains were not significant, and interpretation of intermediate cue as ambiguous was not confirmed |
| **Novak et al., (2015)** | Spatial (Go/No-Go) | 28 female CD-1 | DS+/DS-: Location in an eight-arm radial maze<br><br>AMB: 4 intermediate arms of radial maze (2 near positive, 2 near negative) | Not specified | UCS+: Chocolate flavored pellet<br><br>UCS-: Overhead light | Find a food reward at the end of the POS arms or avoid NEG arms | No DC described | Yes (time spent in positive arm increased, and negative arm decreased across training sessions) | Tail handling (anxiogenic) vs cup handling (non-aversive) | Well documented negative effects of tail handling (increased anxiety) in the literature | No: mice spent more time in ambiguous arms than in POS/NEG arms | No significant handling treatment effect | Less time spent in the AMB arms for tail-handled mice | No | The task failed to detect an effect of handling in the ambiguous cue |
| **Novak et al., (2016a)** | Spatial (Go/No-Go) | 44 female CD-1<br><br>40 female C57BL/6 | DS+/DS-: Location in an eight-arm radial maze<br><br>AMB: 4 intermediate arms of radial maze (2 near positive, 2 near negative) | Not specified | UCS+: Chocolate flavored pellet<br><br>UCS-: Overhead light | Find a food reward at the end of the POS arms and avoid NEG arms | No DC described | Yes (more time spent in positive arm than negative arm) | Varying levels of stereotypic behaviour (SB) | SB occurs in poor-welfare conditions (e.g. prevalent in barren cages)<br><br>However, note: this indicator of affect is prone to false negatives - non-stereotypic animals sometimes have the worst welfare | N/A: responses to AMB cues were not statistically compared the POS and NEG responses | No effect of stereotypy on exploration of ambiguous arms for C57BL/6 mice.<br><br>Some forms of SB predicted more time in ambiguous arms, while others predicted less time in ambiguous arms for CD-1 mice | Less time spent in the AMB arms in stereotypic mice | No | No clear overall effect. But also, this approach is not ideal for construct validation since for identically-housed animals, high SB and low SB individuals do not have clear affective differences[1] |

| Study | Test Type | Subjects | Discriminative stimuli (DS) | DS+ / DS- counterbalanced? | Unconditioned stimuli (UCS) | Trained response | Discrimination criterion (DC) | Successful Discrimination? | Affective state manipulation / inference used in construct validation | Justification / confirmation of affective state manipulation | Were intermediate cues perceived as ambiguous? | Treatment effects on response(s) to ambiguous cues | Required response to ambiguity for successful construct validation | Successful construct validation? (i.e. test is sensitive to affective state) | Reason for failed validation |
|---|---|---|---|---|---|---|---|---|---|---|---|---|---|---|---|
| **Novak et al., (2016b)** | Tactile (Go/Go) | 16 female C57BL/6 /JRcc | DS+/DS-: Grades of sandpaper<br><br>AMB: Intermediate grade sandpaper | Yes | UCS+: Almonds (high-value reward)<br><br>UCS-: No reward<br><br>An oat flake (low-value reward) was always present in an adjacent compartment | Dig for a high-value reward (POS) or low-value reward (NEG) | 10 correct choices (five positive and five negative) out of the 14 free-choice trials per session for two consecutive days | Yes (81% of mice) | Exposure to unpredictable chronic mild stress (UCMS) | Well docuented negative effects of UCMS in literature.<br><br>Assessment saccharine preference (anhedonia) to confirm depressive-like behavior | Control mice showed a graded response, whereas mice exposed to UCMS did not differentiate between near-negative, intermediate, and near-positive cues | No effect of UCMS | Increased AMB latency in UCMS mice | No | No treatment differences in response to ambiguous cues; plus UCMS mice did not show a decreased saccharine preference (suggesting UCMS not very effective here) |
| **Jones et al., (2017) Exp 1** | Auditory (Go-pos/NoGo-neg) | 16 male C57BL/6J | DS+/DS-: 2 kHz at 76 dB or 8 kHz at 65 dB<br><br>AMB: Intermediate auditory tones (3,4,5,6,7 kHz) (from near positive to near negative) | Yes | UCS+: Condensed milk<br><br>UCS-: Air puff | Move to the adjacent compartment to receive a reward (POS) and don't move to avoid punishment (NEG) | 3 training phases: and mice had to achieve 70% accuracy in each | No | No manipulation | N/A | Yes | N/A | N/A | No | No attempt at construct validation (animals also did not learn the task) |
| **Jones et al., (2017) Exp 2** | Auditory (Go-neg/NoGo-pos) | 16 male C57BL/6J | DS+/DS-: 2 kHz at 76 dB or 8 kHz at 65 dB<br><br>AMB: Intermediate auditory tones (3,4,5,6,7 kHz) (from near positive to near negative) | Yes | UCS+: Condensed milk<br><br>UCS-: Air puff | Don't move to receive a reward (POS) or move to the adjacent compartment to avoid punishment (NEG) | 3 training phases: and mice had to achieve 70% accuracy in each | Yes (37% of mice) | No manipulation | N/A | Yes | N/A | N/A | No | No attempt at construct validation |
| **Hintze et al., (2018)** | Spatial (Go/No Go) | 24 female C57BL6/JRj<br><br>24 female RjOrl:SWISS | DS+/DS-: Location of open goal hole at end of arena<br><br>AMB: 3 0pen goal–holes at intermediate locations ( from near positive to near negative) | Yes | UCS+: Chocolate flavored pellet<br><br>UCS-: Nothing | Look for the reward in the goal hole (POS), refrain from responding/or re-initiate a new trial (NEG) | 80% correct responses in positive and negative trials across 4, 20-trial training blocks | Yes (95% of mice) | Housed mice with different types of environmental enrichment<br><br>note: enrichment not compared in statistical analyses | N/A | Yes | N/A | N/A | No | No attempt at construct validation |
| **Bailoo et al., (2018)** | Spatial (Go/No Go) | 96 female C57BL6/JRj<br><br>96 female RjOrl:SWISS | DS+/DS-: Location of open goal hole at end of arena<br><br>AMB: 3 0pen goal–holes at intermediate locations ( from near positive to near negative) | Yes | UCS+: Chocolate flavored pellet<br><br>UCS-: Nothing | Look for the reward in the goal hole (POS), refrain from responding/or re-initiate a new trial (NEG) | 80% correct responses in positive and negative trials across 4, 20-trial training blocks | Yes (98% of mice) | Environmental enrichment | Well documented positive effects of environmental enrichment in literature and observed changes in SB | Yes | No consistent effects of housing on responses to ambiguous cues were found. In the near negative trial C57BL/6 mice showed less Go responses, but the super enriched Swiss mice showed the lowest proportion of Go responses during the same trial (effects were not significant) | Lower proportion of Go responses in barren housed animals, increasing with environmental complexity | No | Significant treatment effects were not detected in the ambiguous trials, and the effects that were observed were not consistently in the predicted direction. |

| Study | Test Type | Subjects | Discriminative stimuli (DS) | DS+ / DS- counterbalanced? | Unconditioned stimuli (UCS) | Trained response | Discrimination criterion (DC) | Successful Discrimination? | Affective state manipulation / inference used in construct validation | Justification / confirmation of affective state manipulation | Were intermediate cues perceived as ambiguous? | Treatment effects on response(s) to ambiguous cues | Required response to ambiguity for successful construct validation | Successful construct validation? (i.e. test is sensitive to affective state) | Reason for failed validation |
|---|---|---|---|---|---|---|---|---|---|---|---|---|---|---|---|
| **Krakenberg et al. (2019a) Exp 1** | Visual (Go/Go) | 6 male C57BL/6J | DS+/DS-: White bar location on tactile screen<br><br>AMB: 3 intermediate locations of the white bar (from near positive to near negative) | No | UCS+: 12 μl condensed milk<br><br>UCS-: 5 sec timeout, lights on<br><br>4 μl of sweet condensed milk (low-value reward) was always delivered when mice touched the adjacent cross | Touch one of the two crosses displayed next to the white bar for a high value reward (POS) or low value reward (NEG) | 5 training phases: requiring 80% correct responses in phases 2-4 | Yes (100% of mice) | No manipulation | N/A | Yes | N/A | N/A | No | No attempt at construct validation |
| **Krakenberg (2019a) Exp 2** | Spatial (Go/Go) | 12 male C57BL/6J | DS+/DS-: Length of tunnel leading to unconditioned stimuli<br><br>AMB: 3 intermediate lengths of the tunnels (from near positive to near negative) | Yes | UCS+: large almond piece<br><br>UCS-: no reward<br><br>A small almond (low-value reward) was always in the adjacent pot during the negative trial | Dig in the correct pot for a large reward (POS) or a small reward (NEG) | 3 training phases: requiring 75% correct responses in phase 1, then 73% correct responses in phase 2 and 3 | Yes (75% of mice) | No manipulation | N/A | Yes | N/A | N/A | No | No attempt at construct validation |
| **Krakenberg et al., (2019b)** | Visual (Go/Go) | 15 male 5HTT (+/+)<br><br>14 male 5HTT (+/-)<br><br>11male 5HTT (-/-) | DS+/DS-: White bar location on tactile screen<br><br>AMB: 3 intermediate locations of the white bar (from near positive to near negative) | No | UCS+: 12 μl condensed milk<br><br>UCS-: 5 sec timeout, lights on<br><br>4 μl of sweet condensed milk (low-value reward) was always delivered when mice touched the adjacent cross | Touch one of the two crosses displayed next to the white bar for a high value reward (POS) or low value reward (NEG) | 5 training phases: requiring 80% correct responses in phases 2-4 | Yes (100% of mice) | Compared serotonin transporter (5-HTT) knockout mice: wild-type 5HTT (+/+)<br><br>heterozygous 5HTT (+/-)<br><br>homozygous 5HTT (-/-) | The 5-HTT knockout mouse is a well-validated model for anxiety. In addition, they corroborated anxiety-like behavior in a battery of tests after the JB assay | Yes | No significant differences were detected between genotypes | Homozygous mice being more likely to press NEG cross for AMB cues than heterozygous and wild-type | No | No treatment effect in response to ambigous cues |
| **Verjat et al., (2020)** | Spatial (Go/No-Go) | 39 male house mice | DS+/DS-: Location in a three-arm radial maze<br><br>AMB: Middle arm between POS and NEG | Yes | UCS+: 10% sugar solution<br><br>UCS-: plain water | Approach and consume the positive/less-positive reward | Individuals had to approach and consume the POS reward with a shorter latency than the NEG (low-value) reward for at least two consecutive days, with a minimum difference of 5 seconds | Yes (64% of mice) | No manipulation, animals that differed in levels of exploratory behaviour were compared<br><br>note: tests of exploration were common anxiety tests, so high exploration reflects low anxiety (e.g. open field test) | Authors used tests of exploration or anxiety-like behaviour | Yes | Exploratory animals responded more pessimistically to ambiguous cues (having longer latencies to approach and consume ambiguous rewards) | High anxiety/low exploratory animals respond more pessimistically to ambiguous cues (having longer latencies to approach and consume ambiguous rewards) | No | Treatment effects were in the opposite direction (i.e. low mood associated with relative optimism)[2] |

| Study | Test Type | Subjects | Discriminative stimuli (DS) | DS+ / DS- counterbalanced? | Unconditioned stimuli (UCS) | Trained response | Discrimination criterion (DC) | Successful Discrimination? | Affective state manipulation / inference used in construct validation | Justification / confirmation of affective state manipulation | Were intermediate cues perceived as ambiguous? | Treatment effects on response(s) to ambiguous cues | Required response to ambiguity for successful construct validation | Successful construct validation? (i.e. test is sensitive to affective state) | Reason for failed validation |
|---|---|---|---|---|---|---|---|---|---|---|---|---|---|---|---|
| **Krakenberg et al., (2020)** | Visual (Go/Go) | 24 male C57BL/6J mice | DS+/DS-: White bar location on tactile screen<br><br>AMB: 3 intermediate locations of the white bar (from near positive to near negative) | No | UCS+: 12 µl condensed milk<br><br>UCS-: 5 sec timeout, lights on<br><br>4 µl of sweet condensed milk (low-value reward) was always delivered when mice touched the adjacent cross | Touch one of the two crosses displayed next to the white bar for a high value reward (POS) or low value reward (NEG) | 6 training phases: requiring 80% correct responses in phases 2-6 | Yes (100%) | Mildly adverse experience: repeated confrontation with a dominant male opponent<br><br>Neutral experience: repeated exposure to a clean cage<br><br>Positive experience: presentation of freshly collected female urine | Mildly adverse experience: losing an aggressive confrontation can increase anxiety-like behaviour<br><br>Positive experience: female pheromones reduce anxiety-like behaviour and aggression in male mice and are suggested to induce positive affect (as they trigger ultrasonic courtship vocalizations) | Yes | Mice responded more 'pessimistically' after the positive experience | Exposure to Mildly Adverse Experience makes mice are more likely to press the NEG cross for AMB cues than Neutral and Positive Experience mice. Exposure to Positive Experience makes mice more likely to respond optimistically, pressing the POS cross for AMB cues more more often than Neutral and Mildly Adverse Experience mice. | No | Treatment effect for the positive experience were in the opposite direction, whereas no treatment effect in response to ambiguous cues was seen for the mildly adverse experience |

[1] Mason GJ, Latham N. 2004. Can't stop, won't stop: is stereotypy a reliable animal welfare indicator?. *Anim.Welf.* **13**, S57-69.

[2] These authors did not aim to validate the task. Instead they aimed to use a task that had not yet been validated to detect differences in affective state associated with exploratory behaviour. However, because the tests used are common tests of mouse anxiety, results could be used to confirm failed construct validation.

**KEY**= POS: positive; NEG: negative; AMB: ambiguous; EXP: experiment; DS: discriminative stimuli; UCS: unconditioned stimuli; JB: judgement bias, DC= discrimination criteria; UCMS= unpredictable chronic mild stress

**Table S2:** Experiment 1 and 2 schedule and protocols

| Phase: | Experimental Design | | |
|---|---|---|---|
| | | Experiment 1 | Experiment 2 |
| All | Affective manipulation | Environmental enrichment | Subcutaneous tumour implantation |
| | JB prediction | CH mice are pessimistic (if task is valid) | Mice with xenografts are pessimistic |
| | High value reward | Banana chip | Almond |
| | Low value reward | Rodent chow | Corn flake |
| | DS+ | Mint or vanilla (counterbalanced) | Vanilla |
| | DS- | Mint or vanilla (counterbalanced) | Mint |
| Digging Training | Digging training schedule | 5 days: 2 Pos trials/day | 4 days: 2 Pos trials/day |
| | Digging trial duration | 5 minutes | 5 minutes |
| Discrimination Training | Discrimination training schedule | 10 days: 4 trials/day | 9 days: 2 trials/day |
| | Digging trial order | Days 1-5: 4 trials /day | |
| | | Trial 1: Pos | |
| | | Trial 2: Neg | Trial 1: Pos |
| | | Trial 3: Pos | Trial 2: Neg |
| | | Trial 4: Neg | |
| | | Days 6-10: 4 trials /day* | |
| | Discrimination trial duration | 5 minutes | 3 minutes |
| Testing for Discrimination Criterion (DC) | Testing schedule | 3-5 days (dependent on time to meet DC): 5 trials / day | 3 days: 3 trials/day |
| | Retesting until mice met DC? | Yes | No |
| | Testing phase order | Trials 1 and 2: Pos or Neg ** | Trial 1: Pos |
| | | Trial 3: test trial | Trial 2: Neg |
| | | Trials 4 and 5: Pos or Neg** | Trial 3: Test trial |
| | Test trial duration | 2 minutes | 1 minute |

<table>
<tr><th colspan="6">Trial Details</th></tr>
<tr><th rowspan="2"></th><th rowspan="2">Trial type</th><th colspan="2">Experiment 1</th><th colspan="2">Experiment 2</th></tr>
<tr><th>Scented Arm</th><th>Unscented Arm</th><th>Scented Arm</th><th>Unscented Arm</th></tr>
<tr><td rowspan="2">Digging and Discrimination Training</td><td>Pos training</td><td>DS+ / Banana</td><td>Water / Chow</td><td>DS+ / Almond</td><td>Water / Corn flake</td></tr>
<tr><td>Neg training</td><td>DS- / No reward</td><td>Water / Chow</td><td>DS- / No reward</td><td>Water / Corn flake</td></tr>
<tr><td rowspan="4">Testing for Discrimination and JB (All conducted with no buried rewards)</td><td>Discrimination criterion (DC)</td><td colspan="4">Mice must dig twice as long in the DS+ pot (Pos test) than DS- pot (Neg test), and dig for at least 3 seconds</td></tr>
<tr><td>Pos test</td><td>DS+</td><td>Water</td><td>DS+</td><td>Water</td></tr>
<tr><td>Neg test</td><td>DS-</td><td>Water</td><td>DS-</td><td>Water</td></tr>
<tr><td>Ambiguous test</td><td>Mixture</td><td>Water</td><td>Mixture</td><td>Water</td></tr>
<tr><td colspan="6">* Trials pseudorandomized so mice always had two Positive (Pos) and two Negative (Neg) trials per day<br>** Trials pseudorandomized so mice always had one Pos and one Neg trial before and after the test trial</td></tr>
</table>

**Table S3:** Ethogram of mouse behaviours (c.f. Harper et al., 2015, Nip et al., 2019) used to confirm relative differences in affective state induced by EH and CH in Experiment 1. Behaviours of interest (stereotypic behaviour and laying inactive but awake) are shaded in grey

| Category | Behaviour | Description |
|---|---|---|
| **Stereotypic Behaviour** | Bar-mouthing | Mouse holds cage bar in diastema and sham-bites for 3 or more seconds |
| | Back flipping | Mouse throws body in backward arch off cage floor or wall and lands in the initial body orientation for 3 or more repetitions |
| | Twirling | Mouse hangs by its forepaws on cage lid and turns body in tight circles for 3 or more repetitions |
| | Patterned lid Climbing | Mouse walks/runs on cage lid using all four paws in a fixed, idiosyncratic route for 3 or more repetitions |
| | Route tracing | Mouse walks/runs on cage floor in a fixed idiosyncratic route, for 3 or more repetitions |
| | Digging | Mouse rapidly digs at bare cage floor with forepaws while mouthing the wall or floor of the cage for 3 or more seconds |
| **Borderline Stereotypic Behaviour** | Borderline twirling | Mouse hangs by its forepaws on cage lid and turns body in tight circles for 1-2 repetitions |
| | Borderline backflipping | Mouse throws body in backward arch off cage floor or wall and lands in the initial body orientation for 1-2 repetitions |
| **Partial Stereotypic Behaviour*** | Partial twirling | Mouse hangs from cage lid with forepaws and turns body in tight circles, while 'catching' the cage lid with her back paws halfway through the turn for 3 or more repetitions |
| | Partial backflipping | Mouse jumps to cage lid from cage floor and hangs there momentarily before falling backward in full rotation of body for 3 or more repetitions |
| | Partial route tracing | Mouse walks/runs on cage floor in no clear pattern and with no clear destination for 3 or more repetitions |
| | Partial Patterned lid climbing | Mouse walks/runs on cage lid using all four paws in no clear pattern and with no clear destination, for 3 or more repetitions |
| **Agonism/Aggression** | Administration | Mouse displaces another mouse from a resource, pushes, grabs, or walks over/under another mouse, chases, pins, mounts or tackles another mouse |
| | Receipt | Mouse receives agonism or aggression as described above |
| **Normal Activity** | General locomotion | All locomotive behaviour performed on the lid and cage floor, except stereotypic behaviour |
| | Maintenance | Nest building, digging at bedding or nesting material, feeding and drinking |
| **Grooming** | Allo-grooming | Mouse grooms another mouse |
| | Receipt of allo-grooming | Mouse is groomed by another mouse |
| | Auto-grooming | Mouse grooms herself |
| | Mixed grooming | Grooming occurs between two or more mice and may alternate between allo-grooming & auto-grooming |
| | Rough allo-grooming/barbering | Mouse pulls the hair or whiskers of another mouse while pushing her paws down on the other mouse. |
| | Receipt of rough allo-grooming/barbering | Mouse is pinned down by another mouse and has her hair or whiskers plucked/pulled, sometimes accompanied by audible vocalizations |
| **Inactive but awake** | | Mouse is motionless, muzzle in sight, and eyes wide open |
| **Inactive** | Sleeping | Lying or sitting curled up with eyes closed |
| | Ambiguous | Mouse is sitting curled up with eyes slightly open, it is unclear whether she is sleeping or not. |
| **Other** | Enrichment use | Using enrichments; excludes stereotyping on them |
| | Wheel running | Mouse runs on wheel |
| | Out of sight inactive | Mouse is still, but out-of-sight and experimenter cannot see the mouse's eyes |
| | Out of sight active | Specific activity is obscured from observer's view, but the mouse is clearly active |
| | Investigation of cage lid | Mouse rears up on hind legs and sniffs at or through the cage bars, occasionally biting at them |
| | Other | Behaviours not listed here |

*Partial stereotypies may also be borderline. E.g. mouse does a partial backflip for less than 3 repetitions (Code: PB BKFL).

**Tables S4:** Descriptive statistics for each experimental group in Experiment 1 (a-d) and Experiment 2 (e-f)

a)

| Experiment 1 Latency (2min) | | | | | | | |
|---|---|---|---|---|---|---|---|
| **Trial** | **DS+** | **Housing** | **N** | **Mean** | **Std Dev** | **Minimum** | **Maximum** |
| Ambiguous | Mint | CH | 10 | 19.85 | 13.8585112 | 3 | 38 |
| | | EH | 5 | 15.9 | 14.7918897 | 2 | 35 |
| | Vanilla | CH | 7 | 65.5714286 | 42.4641305 | 24 | 120 |
| | | EH | 9 | 39.8888889 | 39.3230036 | 3 | 120 |
| Negative | Mint | CH | 10 | 59.3 | 47.6469656 | 7 | 120 |
| | | EH | 5 | 60.9 | 57.6209597 | 4 | 120 |
| | Vanilla | CH | 7 | 107.714286 | 25.4670656 | 52 | 120 |
| | | EH | 9 | 101.944444 | 37.5153672 | 17 | 120 |
| Positive | Mint | CH | 10 | 14.05 | 10.2617142 | 3 | 31 |
| | | EH | 5 | 21.5 | 19.9749844 | 4 | 56 |
| | Vanilla | CH | 7 | 13.5714286 | 10.8144524 | 4 | 36 |
| | | EH | 9 | 15.2222222 | 12.5973983 | 2 | 40 |

b)

| Experiment 1 Digging Time (2min) | | | | | | | |
|---|---|---|---|---|---|---|---|
| **Trial** | **DS+** | **Housing** | **N** | **Mean** | **Std Dev** | **Minimum** | **Maximum** |
| Ambiguous | Mint | CH | 10 | 20.3314 | 14.8314848 | 4 | 50 |
| | | EH | 5 | 11.5257 | 4.8523484 | 3 | 16 |
| | Vanilla | CH | 7 | 2.523 | 2.8093922 | 0 | 8 |
| | | EH | 9 | 3.3411111 | 4.048736 | 0 | 12 |
| Negative | Mint | CH | 10 | 2.0779 | 2.3275353 | 0 | 7 |
| | | EH | 5 | 3.1641 | 3.6364928 | 0 | 9 |
| | Vanilla | CH | 7 | 1.1167857 | 2.067503 | 0 | 5 |
| | | EH | 9 | 0.5485556 | 1.4505334 | 0 | 4 |
| Positive | Mint | CH | 10 | 11.68295 | 7.4318319 | 4 | 30 |
| | | EH | 5 | 11.5133 | 6.860568 | 4 | 20 |
| | Vanilla | CH | 7 | 15.8593571 | 11.5619775 | 10 | 41 |
| | | EH | 9 | 19.7060556 | 12.842045 | 6 | 42 |

c)

| Experiment 1 Latency Time (1min) | | | | | | | |
|---|---|---|---|---|---|---|---|
| **Trial** | **DS+** | **Housing** | **N** | **Mean** | **Std Dev** | **Minimum** | **Maximum** |
| Ambiguous | Mint | CH | 10 | 19.85 | 13.8585112 | 3 | 38 |
| | | EH | 5 | 15.9 | 14.7918897 | 2 | 35 |
| | Vanilla | CH | 7 | 44.7142857 | 15.942232 | 24 | 60 |
| | | EH | 9 | 31.0555556 | 23.7413727 | 3 | 60 |
| Negative | Mint | CH | 10 | 38.15 | 20.7378478 | 7 | 60 |
| | | EH | 5 | 36.9 | 29.2262724 | 4 | 60 |
| | Vanilla | CH | 7 | 58.8571429 | 3.0237158 | 52 | 60 |
| | | EH | 9 | 55.1666667 | 14.5 | 17 | 60 |
| Positive | Mint | CH | 10 | 14.05 | 10.2617142 | 3 | 31 |
| | | EH | 5 | 21.5 | 19.9749844 | 4 | 56 |
| | Vanilla | CH | 7 | 13.5714286 | 10.8144524 | 4 | 36 |
| | | EH | 9 | 15.2222222 | 12.5973983 | 2 | 40 |

d)

| Experiment 1 Digging Time (1min) | | | | | | | |
|---|---|---|---|---|---|---|---|
| **Trial** | **DS+** | **Housing** | **N** | **Mean** | **Std Dev** | **Minimum** | **Maximum** |
| Ambiguous | Mint | CH | 10 | 12.4385 | 11.1114827 | 2 | 38 |
| | | EH | 5 | 8.9175 | 3.4884964 | 3 | 12 |
| | Vanilla | CH | 7 | 1.9612143 | 2.5642692 | 0 | 7 |
| | | EH | 9 | 2.9015 | 3.0879871 | 0 | 9 |
| Negative | Mint | CH | 10 | 1.96365 | 2.6208867 | 0 | 8 |
| | | EH | 5 | 2.506 | 3.2635347 | 0 | 8 |
| | Vanilla | CH | 7 | 0.1678571 | 0.4441083 | 0 | 1 |
| | | EH | 9 | 0.0644444 | 0.1933333 | 0 | 1 |
| Positive | Mint | CH | 10 | 10.6814 | 7.180382 | 6 | 30 |
| | | EH | 5 | 11.1254 | 5.9116844 | 4 | 17 |
| | Vanilla | CH | 7 | 15.3425 | 11.2306047 | 7 | 40 |
| | | EH | 9 | 17.1118333 | 11.6984165 | 6 | 40 |

e)

| Experiment 2 Latency | | | | | | | |
|---|---|---|---|---|---|---|---|
| **Trial** | **Sex** | **Treatment** | **N** | **Mean** | **Std Dev** | **Minimum** | **Maximum** |
| Ambiguous | Female | Control | 5 | 48.00 | 21.68 | 10 | 60 |
| | | Tumour | 7 | 36.29 | 23.68 | 5 | 60 |
| | Male | Control | 8 | 12.00 | 15.89 | 2 | 42 |
| | | Tumour | 9 | 39.78 | 26.27 | 3 | 60 |
| Negative | Female | Control | 5 | 29.60 | 27.79 | 7 | 60 |
| | | Tumour | 7 | 56.14 | 10.21 | 33 | 60 |
| | Male | Control | 8 | 45.13 | 22.17 | 3 | 60 |
| | | Tumour | 9 | 22.33 | 24.53 | 2 | 60 |
| Positive | Female | Control | 5 | 16.00 | 16.76 | 3 | 45 |
| | | Tumour | 7 | 16.14 | 17.80 | 1 | 39 |
| | Male | Control | 8 | 9.38 | 13.87 | 2 | 41 |
| | | Tumour | 9 | 8.00 | 6.02 | 2 | 17 |

f)

| Experiment 2 Digging Time | | | | | | | |
|---|---|---|---|---|---|---|---|
| **Trial** | **Sex** | **Treatment** | **N** | **Mean** | **Std Dev** | **Minimum** | **Maximum** |
| Ambiguous | Female | Control | 5 | 2.60 | 3.71 | 0 | 8 |
| | | Tumour | 7 | 5.14 | 7.80 | 0 | 22 |
| | Male | Control | 8 | 8.50 | 5.21 | 1 | 15 |
| | | Tumour | 9 | 1.44 | 2.19 | 0 | 6 |
| Negative | Female | Control | 5 | 2.80 | 3.03 | 0 | 6 |
| | | Tumour | 7 | 0.57 | 1.51 | 0 | 4 |
| | Male | Control | 8 | 1.75 | 2.19 | 0 | 5 |
| | | Tumour | 9 | 2.56 | 2.30 | 0 | 7 |
| Positive | Female | Control | 5 | 18.00 | 9.17 | 9 | 33 |
| | | Tumour | 7 | 12.71 | 5.41 | 5 | 21 |
| | Male | Control | 8 | 14.00 | 4.84 | 9 | 22 |
| | | Tumour | 9 | 17.78 | 9.68 | 4 | 30 |

**Figure S1:** Total seconds digging during positive, negative and ambiguous test trials in Experiment 2

Total seconds digging during positive, negative and ambiguous test trials. (a) Total digging in female (F, n=12) or male (M, n=17) nude mice (data logarithmically transformed); (b) Total digging male control (C, n=8) and tumour-bearing (T, n=9) mice. Digging duration was not construct validated in Experiment 1, which is why it was not used to test mood-related hypotheses in Experiment 2. Nevertheless, in males, the sex in which cancer status affected 'pessimistic' digging latencies in ambiguous trials, mice with tumours also showed reduced digging durations in these same trials. This suggests that it would be valuable to re assess the construct validity of digging duration, by avoiding counter-balancing to reduce the noise that this reintroduces [2], and/or supplementing the housing manipulation with other treatments designed to increase effects size, e.g. differential handling as in [3]

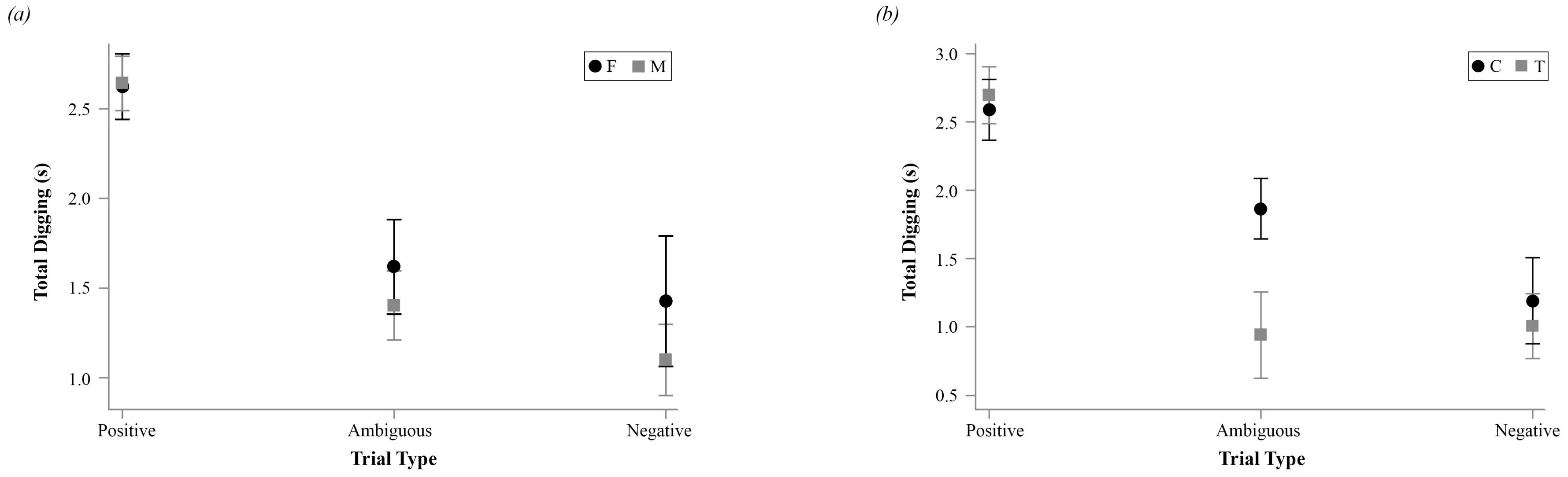


2. Jones S, Neville V, Higgs L, Paul ES, Dayan P, Robinson ESJ, Mendl M. 2018 Assessing animal affect: an automated and self-initiated judgement bias task based on natural investigative behaviour. Sci. Rep. 8, 12400. (doi:10.1038/s41598-018-30571-x)
3. Gouveia K, Hurst JL. 2017 Optimising reliability of mouse performance in behavioural testing: The major role of non-aversive handling. Sci. Rep. 7, 44999. (doi:10.1038/srep44999)

**Tables S5:** ANOVA tables for selected models used for Experiment 2: a) latency, b) digging time

a) Experiment 2 Latency:

| Type III Tests of Fixed Effects | | | | |
|---|---|---|---|---|
| **Effect** | **Num DF** | **Den DF** | **F Value** | **Pr > F** |
| **Trial** | 2 | 58.68 | 15.64 | <.0001 |
| **Treatment** | 1 | 30.12 | 0.20 | 0.6612 |
| **Trial* Treatment** | 2 | 58.68 | 1.08 | 0.3471 |
| **Sex** | 1 | 30.12 | 6.20 | 0.0185 |
| **Sex*Trial** | 2 | 58.68 | 0.81 | 0.4513 |
| **Sex* Treatment** | 1 | 30.12 | 0.07 | 0.7894 |
| **Sex*Trial* Treatment** | 2 | 58.68 | 8.77 | 0.0005 |

b) Experiment 2 Digging:

| Type III Tests of Fixed Effects | | | | |
|---|---|---|---|---|
| **Effect** | **Num DF** | **Den DF** | **F Value** | **Pr > F** |
| **Trial** | 2 | 45.59 | 27.76 | <.0001 |
| **Treatment** | 1 | 36.71 | 1.84 | 0.1827 |
| **Trial* Treatment** | 2 | 45.59 | 1.24 | 0.2997 |
| **Sex** | 1 | 36.71 | 0.71 | 0.4048 |
| **Sex*Trial** | 2 | 45.59 | 0.37 | 0.6899 |
| **Sex* Treatment** | 1 | 36.71 | 0.06 | 0.8128 |
| **Sex*Trial* Treatment** | 2 | 45.59 | 0.86 | 0.4290 |
